# Selection for the bacterial capsule in the absence of biotic and abiotic aggressions depends on growth conditions

**DOI:** 10.1101/2020.04.27.059774

**Authors:** Amandine Buffet, Eduardo P.C. Rocha, Olaya Rendueles

## Abstract

Extracellular capsules protect the cell against both abiotic and biotic stresses such as bacteriophages and the host immune system. Yet, it is unclear if capsules contribute to fitness in the absence of external aggressions, in spite of the cost of production. Here, we enquire if there are conditions favouring the presence of the capsule in *Klebsiella*, where it is a major virulence factor. We shortly experimentally evolved 19 different strains, and show that small changes in growth media have a strong impact on the maintenance of the capsule. Competitions with capsule mutants in nine different strains showed that capsules provide ∼20% of fitness advantage in nutrient-poor conditions, due to faster growth rates and higher yields. In contrast, the capsule was readily lost in nutrient-rich media. The growth environment, as well as the capsule serotype, strongly influenced the role of the capsule in major virulence traits like hypermucoviscosity and biofilm formation. Our data shows that the capsule is selected for in situations lacking abiotic, but also biotic stresses and further supports that the capsule role in virulence may be a by-product of adaptation outside the host, hinting that it may have important roles in bacterial physiology yet to be discovered.

**SIGNIFICANCE:** Bacterial capsules are a wide-spread virulence factor that limits efficacy of antimicrobial therapy. Whereas most studies focus on the role of the capsule in pathogenesis, very few have addressed the conditions under which the capsule is primarily selected for. Here, we show that small changes in growth media have a strong impact in the maintenance of the capsule and the fitness advantage they confer. Our results raise the question whether conditions lacking biotic or abiotic stresses, in addition of selecting for the maintenance of the capsule, can also play a role in selecting for serotype variation. Our results further support that the role of the capsule in virulence may be a by-product of adaptation outside the host, hinting that there may be other functionalities yet to be discovered for it.

## INTRODUCTION

Most environments in which bacteria thrive are complex and can be temporally or spatially heterogeneous, both in their abiotic (pH, nutrients, chemicals…) or biotic compositions (niche invasions, extinction events). This can represent an evolutionary challenge for microbes (Levins 1968, Meyers and Bull 2002). In order to survive, bacterial species can either move to less stressful territories, adapt to the new circumstances or resist until conditions are suitable (Rittershaus et al 2013). Notably, bacteria can produce a thick extracellular layer – the capsule – that has been reported to enhance survival and increase tolerance to external aggressions (Ophir and Gutnick 1994, Tipton et al 2018). Extracellular capsules constitute the first barrier between the cell and its environment and are encoded in half of the bacterial genomes, in some archaea and yeast (Rendueles et al 2017). Capsules fulfill many roles in bacterial physiology. They have been associated with resistance to abiotic stresses such as UV light and desiccation (Ophir and Gutnick 1994, Tipton et al 2018). However, they are mostly studied for their role enhancing survival under biotic stresses, including resistance to grazing protozoa (Jung et al 2007), antibiotics (Fernebro et al 2004, Geisinger et al 2019), antimicrobial peptides (Band and Weiss 2015, Campos et al 2004), and host immune defences, such as human serum and phagocytes (Podschun and Ullmann 1992, Tomas et al 1986, Williams et al 1983).

The majority of capsules belong to the so-called Group I or Wzx/Wzy-dependent capsules (Rendueles et al 2017). These capsules are high molecular weight polysaccharides made up of repeat units of oligosaccharides. The different possible oligosaccharide combinations and residue modifications lead to different serotypes. Serotype diversity, even within-species, is very large. This genetic diversity has been particularly studied in facultative pathogens such as *Streptococcus pneumoniae* (Bentley et al 2006), *Campylobacter jejuni* (Guerry et al 2012), and *Acinetobacter baumanii* (Giguere 2015). In *Klebsiella pneumoniae*, there are at least 77 serologically-defined capsule serotypes (Mori et al 1989, Orskov 1955, Podschun and Ullmann 1998), and more than 130 serotypes identified through comparative genomics (Follador et al 2016, Wyres et al 2016). This serotype diversity results from the faster evolutionary rates of capsular loci compared to those of the rest of the genome. The former has been shown to occur by homologous recombination (Mostowy et al 2017, Wyres et al 2015) and horizontal gene transfer (Croucher et al 2015, McBride et al 2007, Thrane et al 2015). As a result, the same capsular serotype can be found in multiple lineages of *K. pneumoniae* and a monophyletic clade can include distinct serotypes (Wyres et al 2015, Wyres et al 2019). Selection for rapid variation of capsule serotypes is thought to result from biotic stresses, notably because phages and the human adaptive immune system target the capsule in a serotype specific manner, which favours the emergence of novel serotypes (Mostowy and Holt 2018). This process can be accelerated by exposure to vaccines which target specific capsule serotypes (Croucher et al 2013, Guerry et al 2012, Lamb et al 2014).

Many studies have assessed the role of the capsule during pathogenesis, yet little is known about the primary selective forces that underlie the emergence and maintenance of the capsule when bacteria are not under biotic stress. Since capsules are costly to produce, if they are selected mainly to protect bacterial cells from phages, immune systems, or specific stresses, they should be readily lost in the absence of these challenges. Consequently, serial passaging of *Klebsiella* in lithium chloride peptone water, as well as aging of cultures, readily leads to the emergence of non-capsulated clones (Randall 1939), indicating that the capsule can be easily lost.

To further investigate the conditions where the capsule is advantageous for the cell, we study bacteria from the genus Klebsiella. This genus is composed of ubiquitous free-living and host-associated bacteria able to colonize a large range of environments (Bagley 1985, Wyres et al 2020), due to a diverse metabolism (Blin et al 2017) and the presence of a Wzx/Wzy-dependent capsule (Follador et al 2016). *Klebsiella spp*., and more particularly *K. pneumoniae*, are opportunistic pathogens and major multi-drug resistant (MDR) bacteria that cause both community-acquired and hospital-acquired infections in humans (Bengoechea and Pessoa 2019, Caneiras et al 2019, Lee et al 2017). Most isolated *Klebsiella* have a capsule, and the latter is regarded as the major virulence factor of the genus (Paczosa and Mecsas 2016). To test the maintenance of capsule production in several growth conditions, including host-related environments, we selected nineteen *Klebsiella* strains representative of the genetic diversity of the *K. pneumoniae* species complex (Blin et al 2017). These strains have different capsule serotypes (K-locus) and O-antigens (O-locus), and include a closely-related *K. variicola* (strain #24), and two *K. quasipneumoniae* subsp. *similpneumoniae* (strains #44 & #214) (Table S1). We generated a panel of nine capsule mutants to address the precise fitness effects of the capsule in a range of conditions. Finally, we explored the importance of the serotypes and the genetic background of the strain in driving capsule-related phenotypes. The integration of these results revealed that the capsule can be selected in the absence of biotic stresses.

## RESULTS

### The capsule is lost in nutrient rich environments

Early reports (Julianelle 1928, Randall 1939) and our laboratory observations reveal that serial passaging of *Klebsiella* cultures in rich laboratory media (LB) generates spontaneous mutants lacking the capsule (de Sousa et al 2020). This strongly suggests that capsule has a negative impact on fitness in laboratory environments. To understand the fitness cost of capsule in the absence of biotic pressures, we serially passaged 19 different *Klebsiella* strains belonging to six different serotypes (at least three strains per serotype), among which the virulence-associated KL1, KL2 and the emergent KL107 associated to multi-drug resistance. We then measured the natural emergence of non-capsulated mutants after *ca*. 20 generations in five different growth media (see Methods). Artificial sputum medium (ASM) (Fung et al 2010) and artificial urine medium (AUM) (Brooks and Keevil 1997) mimic host-related nutritional conditions. The remaining are typical laboratory media with decreasing levels of nutrients: rich LB medium, minimal medium supplemented with 0.2% of glucose (M02), and minimal medium supplemented with 0.05% of glucose (M005). Beforehand, we verified that the strains were able to produce a capsule in the different growth conditions by direct microscopic observation and by capsule quantification using the uronic acid method (Methods, Figure S1). To initiate the experiment, we grew three independent clones of each strain for 16 hours in LB and then diluted (1:100) into each growth media. All 19 strains were then serially transferred into fresh media every 24 hours for three days. Although each growth media has different carrying capacities, cultures were always diluted to 1:100 and after 24h of growth and each media was saturated with cells in late stationary phase. This ensured that in all growth media, the different populations underwent a similar number of generations (∼ 20).

Non-capsulated mutants emerged after three days, in 111 of the 290 independently evolved populations (38%) (Figure 1). Most ASM (74%) and LB (89%) lineages showed capsule mutants, and these accounted for 85 of the 111 populations with non-capsulated mutants. In contrast, non-capsulated mutants were very rarely observed in the other three growth media (M02, M005, and AUM). These data suggest that it was the low amount of nutrients present in the growth media that drove the maintenance of the capsule, rather than the opposition between host-related environments versus lab environments. Indeed, it was in the nutrient rich media (ASM and LB), where the capsule was readily lost, whereas it was consistently maintained in the three nutrient poor media (M02, M005 and AUM). To further confirm this, we allowed the experiment to run for 30 days in LB and M02 for a KL2 strain (#26). The three replicate populations that evolved in M02 remained capsulated whereas all populations evolving in LB rapidly and irreversibly lost their capsule (Figure S2).

**Figure 1.**
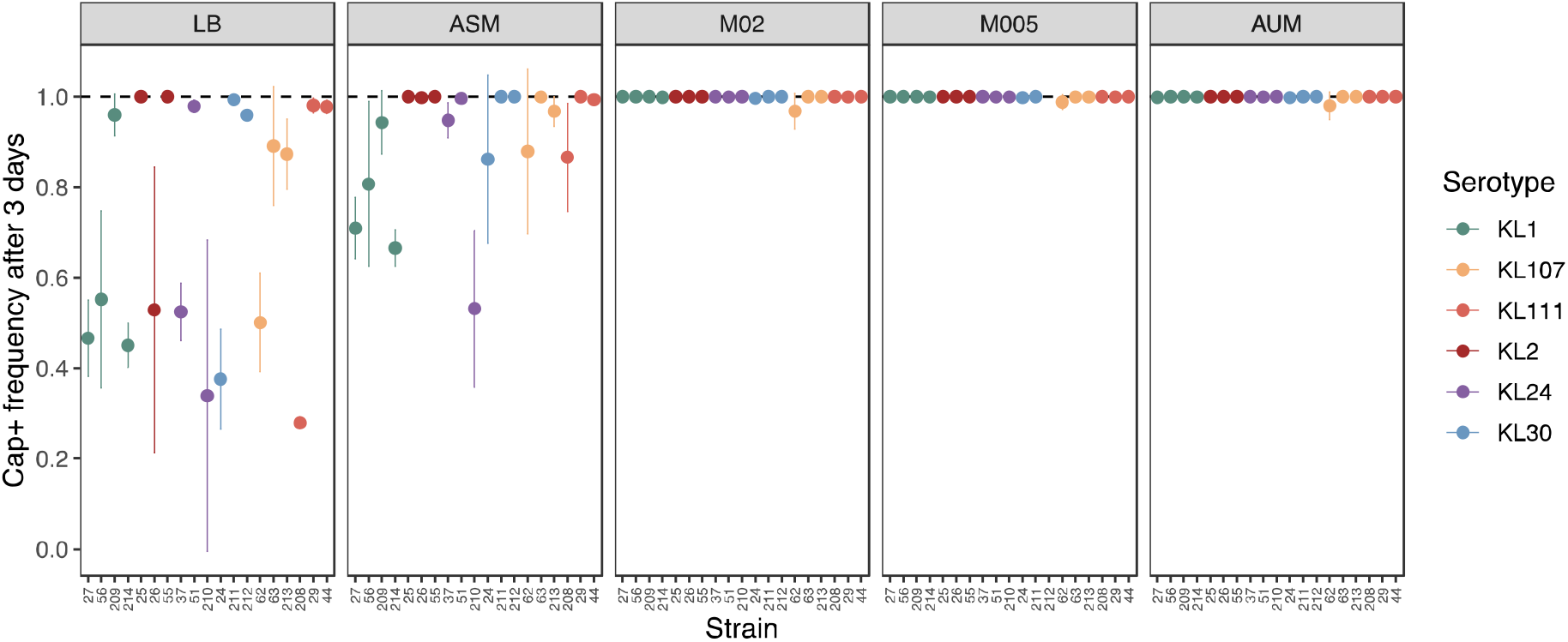
Frequency of capsulated colonies across strains and growth media after three days of evolution. Strains were diluted 1:100 and inoculated in fresh media every 24 hours during three days. Strain #212 was unable to grow in M005. Points represent the average loss of capsule across at least three independently evolving populations per strain. Error bars represent standard deviation across replicate populations.

Some strains seem more prone to lose the capsule. An extreme example concerns strain #62 for which we found capsule mutants in all fifteen independently-evolved populations across the five different growth media. We tested if these differences could be due to the amount of capsule produced by each strain, but the frequency of non-capsulated mutants does not correlate with the amount of capsule produced by each strain before the evolution experiments (P > 0.05, Spearman’s rank correlation). We tested the relevance of both serotype and growth media on the emergence of capsule mutants using a generalized linear model (R^2^ = 0.39). This revealed that the emergence of capsule mutants was both dependent on the growth media (P < 0.001) and on the serotype (P<0.02). This suggests that the capsule is deleterious under rich media and selected for in poor media, a factor that depends at a lesser extent on the capsule serotype.

### The capsule provides a fitness advantage in nutrient poor environments

To precisely assess the costs and benefits of the capsule across growth media, we sought to perform competition experiments between capsulated (Cap+) and their respective isogenic non-capsulated (Cap-) mutants. We thus undertook the construction of in-frame and markerless deletions of *wza*, the outer membrane exporter of the capsule, a well-annotated and essential core gene of the capsule operon. We used the allelic recombination strategy and aimed at deleting *wza* in 15 different strains from different serotypes, for which we had easily obtained *rcsB* in-frame deletion mutants. (RcsB is a capsule regulator and mutants have a reduced capsule production (Dorman et al 2018, Wacharotayankun et al 1992)). We obtained *wza* deletion mutants for five strains, from several serotypes (see Table S1). From these double-recombination events, we isolated both a mutant and a wild type. This ensures that the phenotype was directly due to the *wza* deletion and not to other off-target mutations that could have accumulated in the genome during the mutant generation, and thus present in both the mutant and the wild type. All experiments were performed in parallel with the parental strain, the *Δwza* mutant and the associated wild type from the same double-recombination event. After several attempts, we obtained *wza* deletion mutants in four additional strains, but Illumina sequencing revealed off-target compensatory mutations only present in the mutant (and not in the associated wild type) (see Methods). In the main text, we present statistics and figures for all strains, and in supplemental material, for the five clean mutants. Microscopic observation and uronic acid quantification confirmed that all nine *Δwza* mutants lack a capsule (Figure S3). For the remaining seven strains, no *Δwza* mutants were obtained despite several attempts.

To assess the competitive indexes, or fitness, of the mutant strains, we performed direct competition experiments between wild type strains and their respective *Δwza* mutant strains. The strains were initially mixed in a 1:1 ratio and allowed to compete in all five growth media for a 24-hour period. We estimated that the natural emergence of capsule mutants in rich media is rare during the first 24 hours and is thus negligible in the interpretation of our results. Competition experiments showed a significant effect of the environment in the fitness of the Cap + strains (Figure 2, Kruskal-Wallis, P = 0.0046, N=8). In this analysis, we excluded strain #55 due to a growth deficiency of the *Δwza* mutant. Overall, we observe a marginal decrease of Cap+ genotypes in nutrient rich media (P= 0.07 and P = 0.05 in LB and ASM, respectively, one-sided t-test, difference of 1). Lower fitness in these growth media was significant in 5 out of 8 strains in LB and 4 out of 8 in ASM strains.

**Figure 2.**
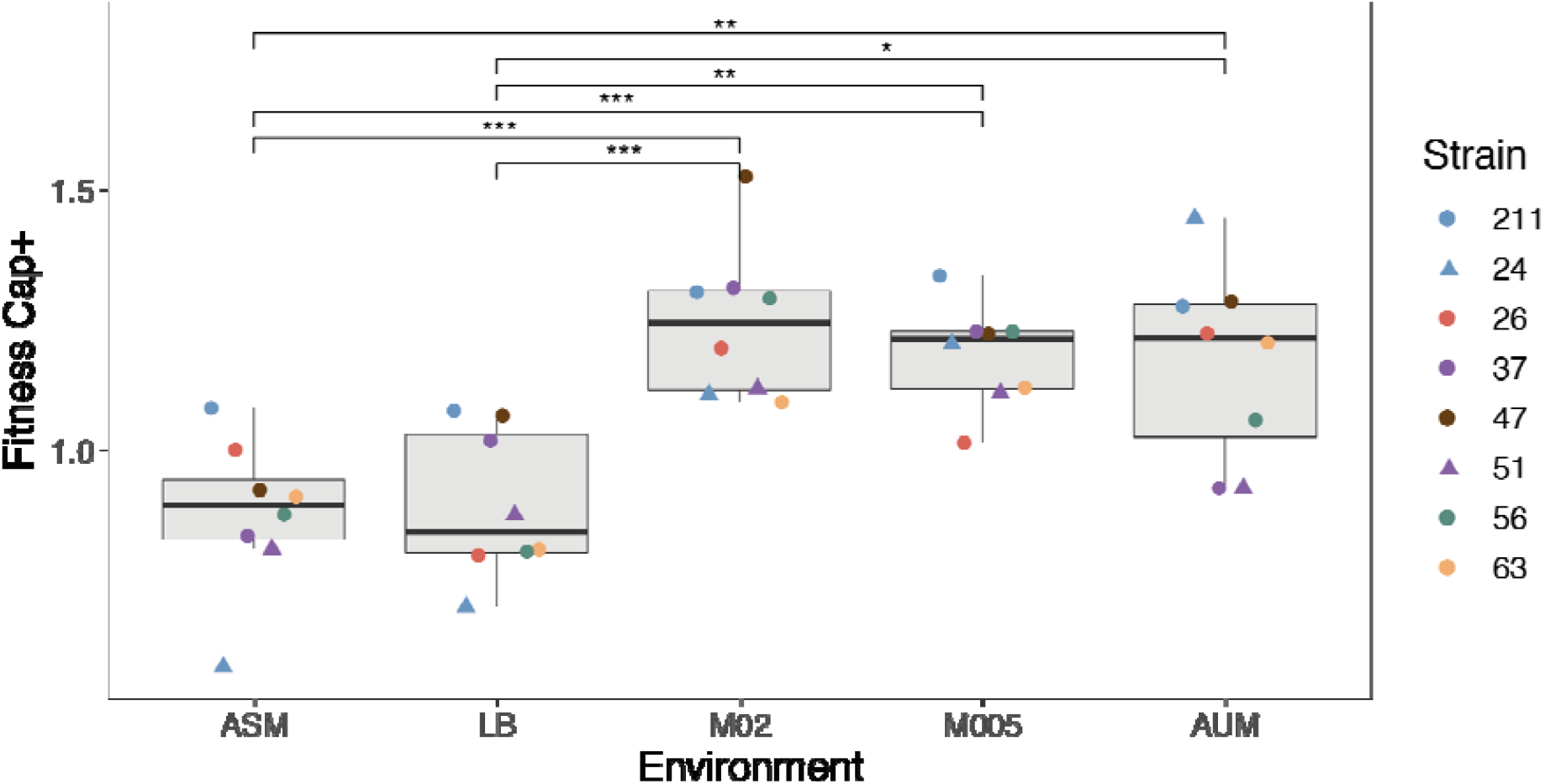
Competition between capsule mutants and their associated wild type in different growth media. Each point represents the average of at least three independent replicates. Individual error bars for each strain are not displayed for clarity purposes. Colours represent different serotypes (as in Figure 1), whereas the shape represent different species. Results for the strains with no compensatory mutations are presented in the supplemental material (Figure S4). * P < 0.05 ; ** P< 0.01, Tukey post hoc and correction for multiple tests.

Opposite to what is observed in nutrient rich media, Cap+ strains have a strong fitness advantage in M02, M05 and AUM (P< 0.05, *N*=8, one-sided t-test, difference of 1). This is in agreement with our previous results that showed the maintenance of the capsule in nutrient poor media. The average fitness advantage provided by the capsule in nutrient poor media is *ca*. 20%. Individually, most strains had a significant fitness advantage in nutrient poor environments, with the exception of the two KL24. In these two strains, the capsule significantly imposed a fitness cost in AUM, suggesting a particular negative interaction between KL24 capsule and AUM (Figure 2). To further corroborate the hypothesis that the amount of nutrients in the media determines the fitness of capsulated strains, we competed the three strains for which the capsule was significantly costly in LB (strain #56, #26 and #24, belonging to three different serotypes) in diluted LB (1:10). Overall, we observe a significant fitness increase of capsulated strains of ∼23% in the diluted rich media (Wilcoxon rank sum test, P< 0.01 for #26 and #24). Qualitative similar results are obtained in diluted ASM (Wilcoxon rank sum test, for strain #24, P< 0.01). Finally, to control for potential fitness side-effects of *wza* deletion, due to accumulation of capsule polysaccharide in the periplasmic space (Rendueles 2020, Tan et al 2020b), we deleted either the *wbaP* or *wcaJ* gene, the first glycosyltransferase of the capsule synthesis pathway, in the abovementioned three strains. Our results consistently show a strong fitness disadvantage of the capsule in rich media, whereas, in nutrient-poor media the capsule provides a significant advantage (Figure S4B, Kruskal-Wallis, P = 0.005). Taken together, our results confirm that the capsule is disadvantageous in rich media, but advantageous in nutrient-poor media.

### The capsule increases growth rate and population yield in nutrient poor growth media

We investigated how the capsule could increase fitness in nutrient poor environments by assessing the effect of mutants in a number of conditions. We first tested whether the capsule provided an advantage during starvation, by increasing cell survival, as shown in *Streptococcus pneumoniae* (Hamaguchi et al 2018). We therefore followed the survival of capsule mutants and wild type under starvation conditions. We allowed cells to starve for two months and counted surviving CFUs every month. As expected, mortality rates of *Klebsiella* are very low compared to other Enterobacteria (Baker et al 2019). Yet, we observed no significant differences between wild type and the capsule mutants after 30 or 60 days (Figure 3A and Figure S5A). To further check if there is a potential effect of the *Klebsiella* capsule on survival, we also challenged the strains to desiccation. No effect of the capsule on cellular survival was observed (*P* > 0.05, t test for significant difference between mutants and wild type, Figure S5B).

**Figure 3.**
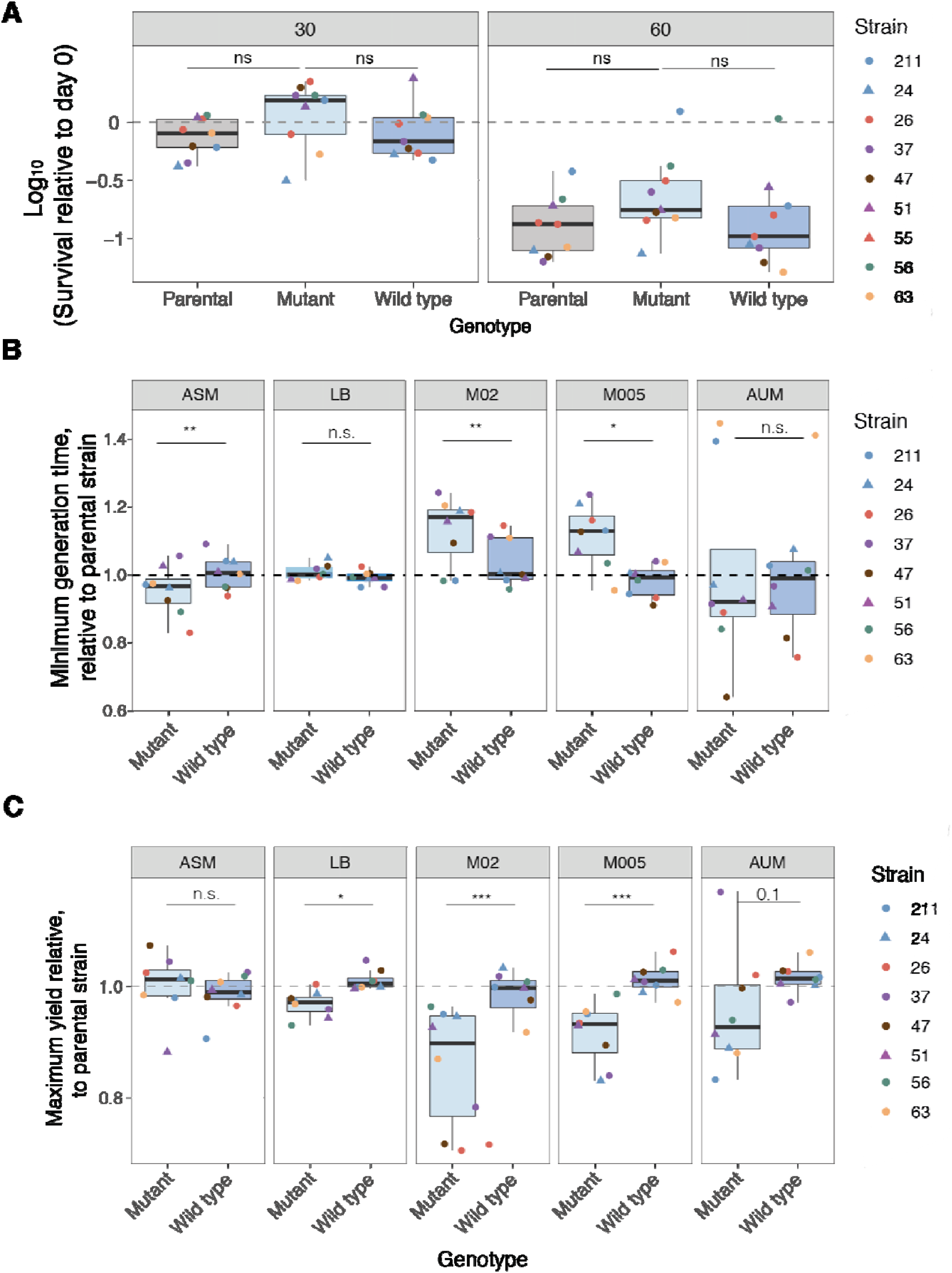
Survival, growth rate and yield of strains in each environment. **A**. Survival of parental, *Δwza* mutant and the associated wild type, relative to day 0. Data is log_10_ - transformed. Individual error bars for each strain are not displayed for clarity purposes. Colours represent different serotypes (as in Figure 1). Different shapes are used to distinguish strains from the same serotype. **B**. Minimum generation time of the mutant and the wild type strains relative to the parental strain in each environment. **C**. Maximum yield of the mutant and the wild type strains relative to the parental strain in each environment. Statistics were calculated using a paired t-test. * P < 0.05.

Given that we observed no significant effect of the presence of the capsule on cellular survival, we tested whether differences in growth rate could account for the fitness effects observed. We thus allowed all strains and their mutants to grow in microtiter plates in all growth media and calculated the minimum generation time (Figure 3B and Figure S6A). As expected, we observed significant differences in growth rate across strains and environments (GLM, P< 0.001 for both, R^2^= 0.60). To assess whether the capsule affects growth dynamics we calculated three different parameters: minimum generation time, maximum yield and the area under the curve (AUC), which takes into account the initial population size, growth rate and carrying capacity of the environment. As expected by the competition experiments, in ASM, we observe that the capsulated strains have slower growth rate and achieve lower absorbances (ODs) a proxy for population yield (Figure 3B and Figure S6A). Interestingly, this is not the case in LB, in which strains reached on average lower yields compared to ASM. In nutrient-poor growth media, we observe the opposite to ASM, namely, that the growth rate of capsulated strains is higher and the populations reach higher yields (Figure 3B and Figure S6B). Control experiments showed a strong correlation between absorbance and cell numbers. Furthermore, direct plating of cells after 16 hours of growth in microtiter plates also revealed larger population sizes in capsulated strains than in mutants in nutrient-poor environments. These observations are qualitatively similar to those observed using AUC calculations. In nutrient-poor environments, notably M02 and M005, the AUC is significantly higher in capsulated strains compared to non-capsulated strains (Figure S6C). Taken together, there is a statistically significant difference in growth rate and yield between the mutants and the wild type in nutrient poor media, suggesting that the fitness benefits of capsulated strains observed in competition are due to advantages in growth dynamics. Qualitatively similar results were obtained using the three *ΔwcaJ* and *ΔwbaP* mutants, namely, a higher yield and AUC of capsulated strains in nutrient poor environments but not in ASM or LB.

We further characterized how the capsule may affect growth to overcome the cost of its production and become an advantage in nutrient poor environments. We first tested if non-capsulated strains exhibit higher death rates in our experimental conditions. The determination of the number of dead cells after 16 hours of growth using a Live-Dead stain suggests this is not so (Two-way ANOVA, P = 0.47, Figure S6D). We then hypothesized that capsulated bacteria could use their capsule as a nutrient source in nutrient poor growth media. To test this, we used the *ΔwcaJ* and *ΔwbaP* mutants, to avoid capsule build up in the periplasm that could interfere with our results. We allowed the mutant strains to grow for 24 and 48 hours in minimal media with no source of carbon, in the presence of purified capsule added exogenously and of uronic acid used as a standard to quantify capsule production (Figure S7). We did not observe significant differences between the treatments where the capsule was added exogenously and the respective controls (Kruskal-Wallis, P > 0.05, Figure S7). These results suggest that the advantage provided by the capsule relies in increased growth rate and yield but not in reduced death rate nor on the ability of the cell to consume the capsule.

### Capsule mutants undergo different fates in host-mimicking media

Our previous results show that changes in growth conditions can strongly affect selection for the capsule. We thus sought to further understand how the capsule affected other traits that are known to be relevant during pathogenesis, and whether this effect is conserved across *Klebsiella* strains. We first addressed the ability of non-capsulated mutants to resist to human serum. Three Cap-mutants were significantly more sensitive to human serum than their Cap+ counterpart, whereas for the other two strains there was no significant difference (Figure 4A). Resistance to human serum can also be due to the presence of different O-antigens (Doorduijn et al 2016, Merino et al 1992). In our panel, only two strains #211 and #26 share the same O-antigen (O1v1, Table S1). These strains have different resistance profiles, suggesting that O-antigen is not solely responsible for serum resistance. These results strongly suggest that the effect of the capsule on resistance to serum is strain-dependent (Multifactorial ANOVA, P < 0.001).

**Figure 4.**
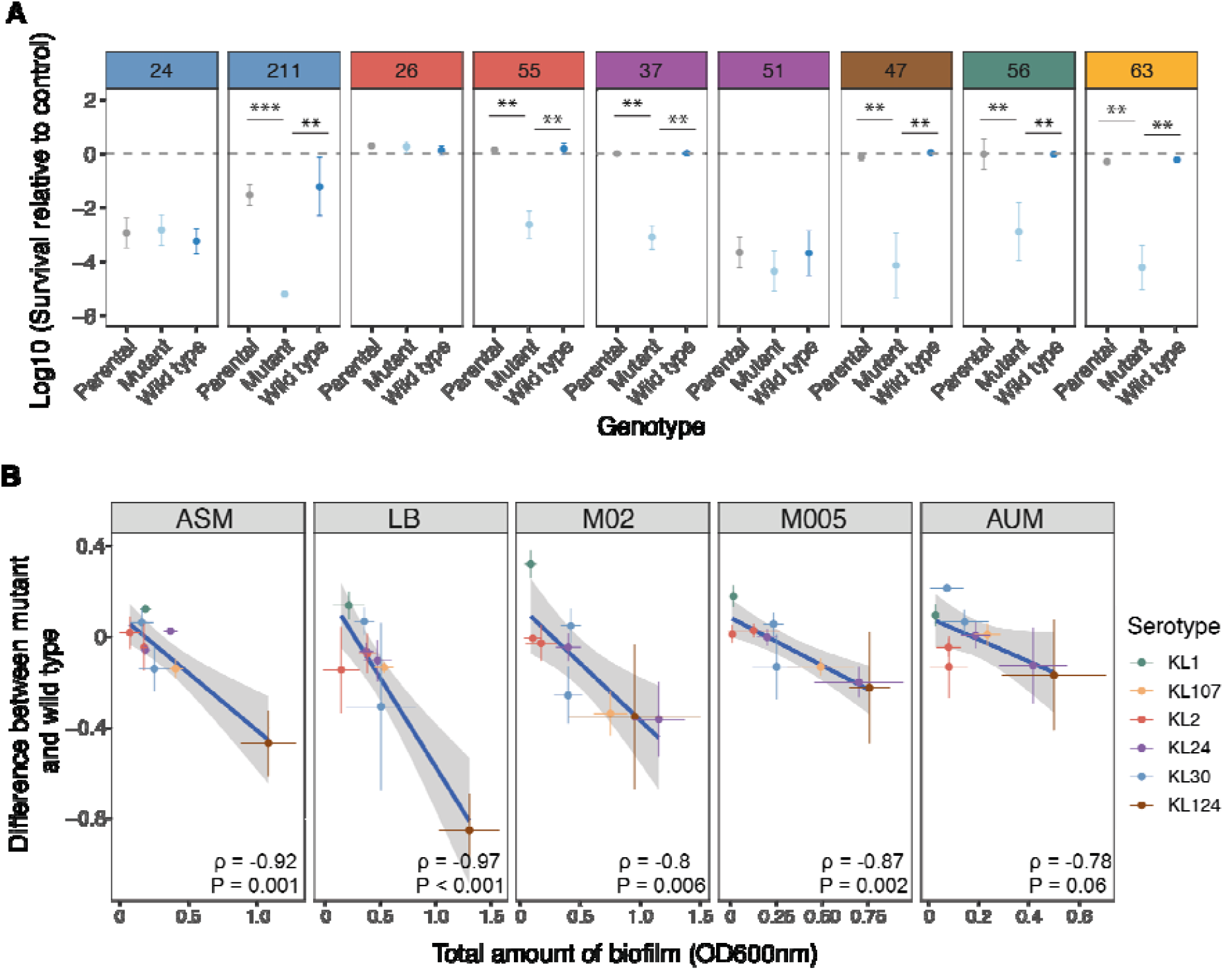
Fate of capsule mutants in host-related context. **A**. Resistance to human serum. Data is presented as relative survival to control (heat-inactivated human serum). * P< 0.05; ** P< 0.01; *** P<0.001 paired t-test. Only significant statistics are represented. **B**. Correlation between the total amount of biofilm formed by the wild type and the difference in biofilm formation between the wild type and the deletion mutant across environments. P-values correspond to Spearman’s rank correlation. Results are qualitatively similar when only taking into account the five strains with no off-target mutations. Absolute values of biofilm formation are shown in Figure S9.

We then explored the ability of capsule mutants to survive in the presence of primary (sodium cholate –CHO-) and secondary (sodium deoxycholate –DCH-) bile salts, since they are antibacterial compounds that may disrupt bacterial cell membranes (Urdaneta and Casadesus 2017). At physiological concentrations (0.05%), we observed no significant sensitivity of *Klebsiella* to bile salts (Figure S8B). When treated with 10X physiological concentrations, some, but not all, capsule mutants showed reduced viability (Figure S8).

Finally, the ability of niche colonization, specially within a host, is directly linked to the ability of the strains to adhere to surfaces and proliferate, that is, to form a biofilm. Cells within biofilms are more resilient, tolerant to antibiotics and less exposed to the host immune system (Hall-Stoodley et al 2004). We thus assessed the ability of the strains to form biofilms and determine how the capsule could impact this trait. It has been previously shown that in some strains of *Klebsiella*, capsule production masks the necessary fimbriae required to attach to the host mucosa (Schembri et al 2004, Schembri et al 2005). On average, we do not observe a difference in the amount of biofilm between the capsulated and non-capsulated mutant forms (P > 0.05, *N* = 9, paired t-test, in all different growth media). However, the amount of biofilm formation in the parental strain is negatively correlated with the difference in biofilm formation between the capsule mutant and its wild type (Figure 4B). In other words, the capsule tends to contribute positively to biofilm formation in strains forming more biofilm. At the strain level, we observe that some non-capsulated mutants adhere significantly more across all environments (strain #56) (Figure S9), whereas other strains have a stronger ability to form biofilm when the capsule is present (strain #47) (Figure S9). This might depend on the capsule composition or it may also result from the absence of adhesion factors in the strain. Overall, our data suggest that it is not the presence or absence of the capsule that affects the ability to form biofilms, but rather the amount of capsule expressed by a given strain.

Taken together, these results show that some properties like resistance to bile salts or human serum, are not inherent to the presence of the capsule, but are rather strain-dependent, or dependent on the serotype-genome interaction.

### Complex interactions between the environment, genetic background and serotype

To further understand the interactions between the serotype, genetic background and environment, we used the original dataset of 19 strains, including at least three from each serotype. We analyzed different capsule-related variables in all five growth media. In addition to the amount of capsule production (Figure S2), the natural emergence of non-capsulated mutants (Figure 1), maximum growth rate, the maximum yield and biofilm formation, we also measured the hypermucoviscosity of strains, a trait linked to hypervirulence and driven, at least partly by the amount of capsule (Lin et al 2012) (Figures S10-S12). Firstly, we used Principal Components Analysis (PCA) to analyze how the different factors (media, serotype or strain) explain the observed variance in the data. The first and second principal components explained more than 60% of the variation observed (Figure 5A). We observed no clear grouping of serotypes according to the two first axes. This suggests that the serotype is not a major determinant of the capsule-related phenotypes (Figure S13). The ellipses regrouping the points by type of media show a small overlap between both rich media (ASM and LB) and these are clearly separated from the two poorest media (M005 and AUM). This suggests that most of the variance is explained by the media, presumably by the amount of nutrients available (Figure 5A). We then used multifactorial ANOVA to analyze quantitatively the effect of the growth media and the serotype on each trait. As expected, we observed a significant effect of the growth media in all measured traits, except in biofilm formation (R^2^ = 0.57, P = 0.06, Table S2). As suggested by the PCA, the influence of the serotype in the capsule-associated traits is less important. Interestingly, the terms of interaction between the environment and the serotype are not statistically significant, suggesting that they are independent (Table S2).

**Figure 5.**
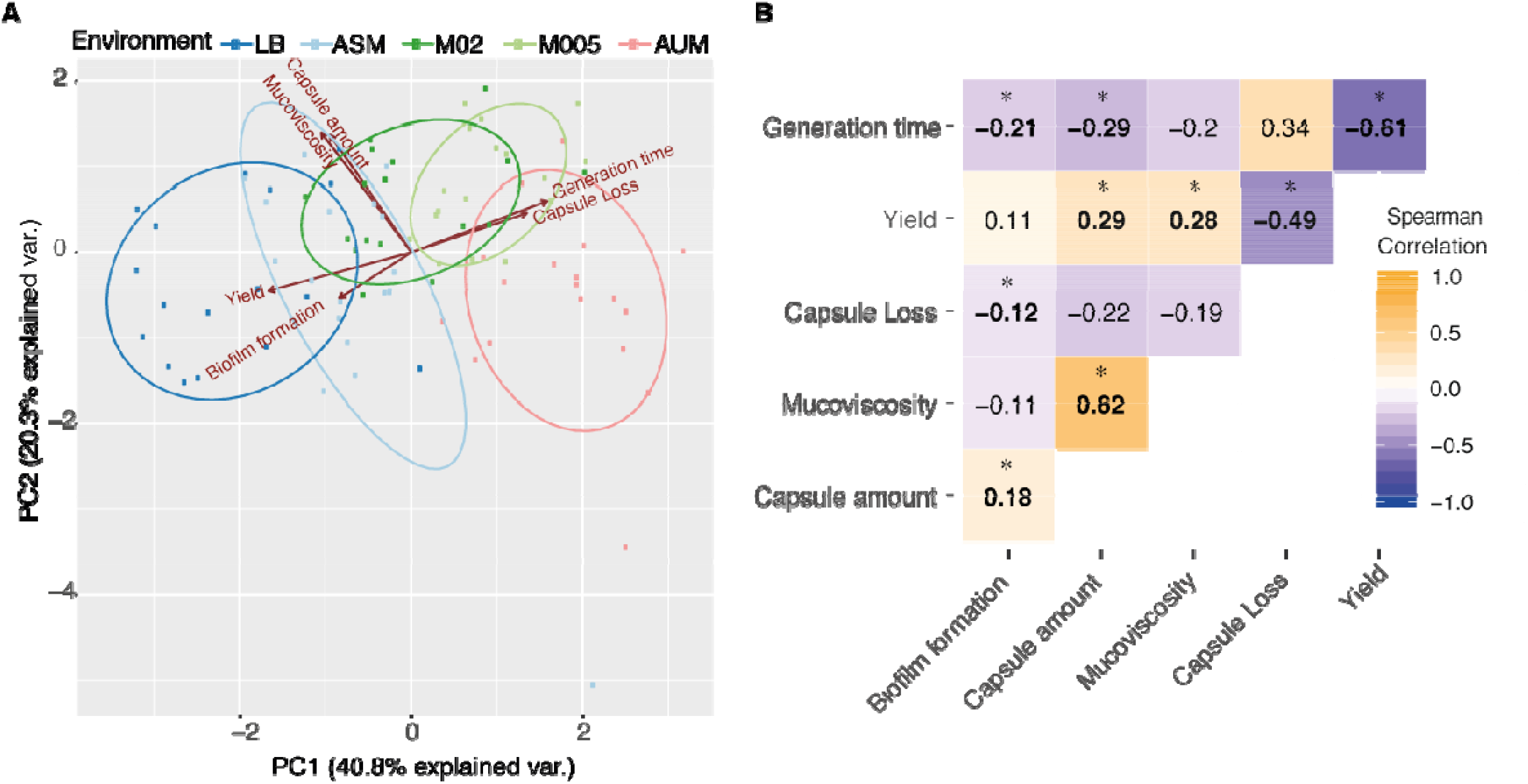
The environment, not the serotype, explains most of the variance observed across capsule-associated traits. **A**. Principal component analyses (PCA) of capsule-related phenotypes. Phenotypes are coloured by environment. Each individual point represents a strain. PCA was performed using the *prcomp* function (options scale and center = TRUE) in R and we used the ggbiplot package for visualization. **B**. Correlation matrix of all traits. Numbers indicate Spearman’s correlation. Stars identify significant correlations (P<0.05).

Secondly, we sought to determine how the different capsule-related variables correlated with each other. The PCA analysis already suggested the existence of such correlations (Figure 5B). As expected, we observe a strong correlation between hypermucoviscosity (HMV) and the amount of capsule, in all individual growth media except in M005. Consistently with previous reports (Lipson 2015, Pfeiffer et al 2001), we observe that the population yield is inversely proportional to the growth rate (Figure 5B). This is significant in all growth media, except M005 (for which *P* = 0.07). Further, the amount of capsule is positively correlated with the maximum yield (Figure 5B), supporting our observation that the capsule contributes to increased yield of the population. In contrast, there is a negative correlation between the amount of capsule and the generation time, a result that is driven by growth in nutrient poor environments (Figure 5B). More precisely, in poor media, bacteria with more capsule are able to grow faster that those producing less capsule. This was not evident from the results of the analyses involving the mutant, as these showed no significant differences in generation time between the capsulated and non-capsulated strains. Overall, our results show that the capsule may impact several important cellular traits, like mucoviscosity and growth, but this is strongly dependent on the environment.

## DISCUSSION

The capsule is an extracellular structure commonly associated to the ability to resist biotic or abiotic aggressions, like the immune system, phages, or desiccation (Bengoechea and Sa Pessoa 2019, Campos et al 2004, Paczosa and Mecsas 2016). In our previous work, we showed that capsules were more prevalent in free-living environmental bacteria than in pathogenic bacteria. This suggested that the role of the capsule in virulence may be a by-product of adaptation outside the host, raising the question of its primary function in cells (Rendueles et al 2017). Here, we explore the conditions in which the capsule is advantageous despite the cost of its production and lack of antagonistic interactions and abiotic aggressions. The evolution experiments and the direct competitions show that the capsule provides a disadvantage during growth in rich medium and an advantage during growth in nutrient poor environments, *ca*. 20%, independently of the capsule mutants used, either outer membrane exporter *Δwza* or the initial glycosyltransferase mutants, *ΔwcaJ* or *ΔwbaP*.

To explain the results of our competition experiments, we examined the effect of the capsule on growth dynamics of wild type strains relative to their capsule mutants. In ASM, the media allowing the highest carrying capacity, we observe that the capsule is costly since mutants grow faster and achieve higher yields. In LB, the results are more complex. Non-capsular mutants readily emerge in the evolution experiments. But there are no significant differences in terms of growth rate between *Δwza* and the wild types, as previously observed in *K. pneumoniae* KPPR1, a strain not included in our panel (Mike et al 2020), and *Acinetobacter baumanii* (Russo et al 2010). We however observe an increased yield and a larger area under the curve in non-capsulated strains, indicative of the cost of the capsule in LB. Similarly, when using the *ΔwcaJ* or *ΔwbaP* mutants, all three measures - AUC, maximum yield and growth rate-show that in LB, the capsule is costly. This might suggest that higher fitness of non-capsulated mutants in competition experiments in LB are caused by higher growth rates during exponential phase. We hypothesize that the differences in growth dynamics between *Δwza* and *ΔwcaJ* mutants are due to the periplasmic concentration of capsule percursors (Rendueles 2020, Tan et al 2020b). Overall, we propose that due to the absence of stresses, capsule inactivation or loss in rich media leads to gains in terms of growth rate and yield, as indicated by the evolution experiments, the competitions with the *Δwza* and *ΔwcaJ* mutants.

In nutrient poor environments, the capsule increases growth rate and populations reach higher yields. This phenomenon explains how the capsule provides a competitive advantage in nutrient poor media, despite its cost in nutrient rich media. Further, we observe a negative correlation between the amount of capsule and generation time in nutrient poor environments (Figure 5B). Thus, bacteria producing more capsule are able to grow faster in nutrient poor environments. These results are somewhat counter-intuitive, as in media in which resources are scarce, sugars should be preserved for essential metabolism including energy-yielding pathways like TCA-cycle, to assemble cellular structures like the cell membrane.

Our results challenge the current view that the primary role of the capsule is to withstand aggressions, because even during growth in pure, stable conditions, the capsule provides a significant advantage and is enough to select for its maintenance. To understand how the capsule could be advantageous for growth in nutrient poor environments, we first hypothesized that capsulated strains could have reduced death rate. We did not observe any effect of the capsule in death rate during the 16 hours of our growth assay. Similarly, our two-month long starvation experiments revealed no differences in survival between capsulated and non-capsulated strains. Previous reports had already shown that in the long term, *Klebsiella spp* are very resilient and have very low death rates (Bossolan et al 2005, Lappin-Scott et al 1988), including in lake water at temperatures close to 0 (Niemela and Vaatanen 1982) or during long-term starvation in PBS as a member of an oral community (Baker et al 2019). Alternatively, capsulated bacteria could use the capsules’ polysaccharides to fuel faster growth and further multiply after consumption of all the environmental resources, as previously reported in *S. pneumoniae* (Hamaguchi et al 2018). However, this is inconsistent with our observation that addition of exogenous capsule to non-capsulated mutants failed to result in a significant increase of CFU in a population. Taken together, and despite of the production costs of the capsule, linked to the biosynthetic process, the energy required to polymerize and export the capsule and the use of resources that are no longer available for other cellular processes, our analysis do not provide a clear explanation on how the capsules increase population yield or growth rate. This will be the object of future research.

Since our results suggests that capsules can be advantageous in the absence of challenges, we re-assessed its impact in features known to be implicated in pathogenesis. Our results show that the impact is very variable across strains and serotypes. We observe an overall strong positive correlation between the capsule and hypermucoviscosity (HMV), a proxy of hypervirulence. This would support the hypothesis that HMV is a result of capsule overproduction. However, not all strains with large capsule are HMV. This is supported by recent studies performed in the KPPR1 strain that show that capsule overproduction is not required for HMV (Mike et al 2020, Walker et al 2019). Moreover, we also show that the capsule serotype affects HMV (P = 0.002), suggesting that the role of the capsule in HMV is convoluted, dependent on the serotype and/or the strain (Walker and Miller 2020).

Similar to the HMV results, we observe that the role capsule in resistance to human serum is also not conserved across strains. Our results are in agreement with a previous study in which cells from different serotypes were first exposed to depolymerases, to digest the capsule (Majkowska-Skrobek et al 2018), and then tested for survival after incubation with human serum and phagocytosis. Consistently, some non-capsulated strains were more sensitive to human serum, but others seem to be as resistant as the control cultures to which no depolymerase was previously added. The diverse impact of the capsule in serum resistance is also supported by a recent study which revealed that most genes contributing to serum resistance were strain-specific (Short et al 2020). This suggests that there may be strong epistasy at play between the capsule and the rest of the genome leading to full serum resistance.

Finally, the strains were naturally resistant to bile salts, suggesting no protective role for the capsule from bile salts at physiological levels. When bile salt concentrations are increased by a factor of ten, the capsule significantly increases survival in some strains but not all. Our results highlight that the role of capsules in traits that are relevant during pathogenesis are diverse and may depend on the strains, the environment, the serotype, and genetic interactions with the background. This implies that the role of capsules in pathogenesis varies across *Klebsiella*. This is supported by the fact that some serotypes are associated to hypervirulence, like K1 and K2, whereas other serotypes are less virulent but associated to multi-drug resistant like KL107 (Wyres et al 2019). In an experiment in which the capsule locus was exchanged between strains with a K2 and K21a serotype, it was shown that the capacity to escape from macrophages, and thus virulence, was inherited with the capsule serotype (Ofek et al 1993). The fact that the roles of the capsule during pathogenesis are not conserved also supports the notion that the benefits of the capsule for virulence are a by-product of adaptation in non-host related contexts. Accordingly, a recent study showed that carbapenem-resistant *K. pneumoniae* ST258 unencapsulated isolates formed more biofilm and could invade the epithelial bladder tissue more effectively than capsulated isolates. This resulted in the formation of intracellular infective reservoirs that lead to recurrent urinary infections (Ernst et al 2020). Overall, our analyses of multiple strains with different serotypes revealed that the role of capsule in traits that may be relevant during pathogenesis is complex and depends not only on the strain but also the serotype.

To conclude, the complex relation between the capsule and the host genome highlights that there are yet many research venues to be explored in terms of selection forces acting on the maintenance and diversification of the capsule, its function across a diversity of strains, and potential epistasy between the capsule and the rest of the genome. Here, we show that the growth media affects the relative fitness of capsulated bacteria, and alone justifies the maintenance of the capsule in a population. Most literature suggests that the rapid turnover of capsular serotypes is due to biotic pressures, such as phage predation, cell-to-cell interactions, the host adaptive immune system, or protozoa grazing (Cobey and Lipsitch 2012, Mostowy and Holt 2018), but more work is needed to elucidate whether the abiotic conditions of the environment also play a role in the diversification of the capsule composition. For example, different capsules could be better adapted to certain poor environments because of their different propensity to aggregate or because of their organization at the cell surface (Phanphak et al 2019, Rendueles 2020). It could also be that different serotypes are associated to specific functions or fitness advantages (Ofek et al 1993). This might lead to selection of serotype switching when bacteria colonize novel environments. Finally, more work is needed to elucidate capsule functions other than those that are commonly evoked and provide more insights into the non-universality of the role of capsules both within and across species (Rendueles 2020). Indeed, recent studies show that the capsule does not protect against temperate or virulent phages in *K. pneumoniae*, but rather the opposite, the capsule acts as a receptor for phages (de Sousa et al 2020, Tan et al 2020a). Taken together, our work suggests that the primary evolutionary forces selecting for the existence of a capsule may be very different from those selecting for its subsequent diversification, and highlights that much work is still needed to fully understand the primary role of capsules, both at the molecular and at the ecological level.

## Supporting information

Mat Sup

## MATERIALS AND METHODS

### Bacterial strains and growth conditions

Bacterial strains and plasmids used in this study are listed in Table S1. Primers used for the construction of mutants are listed in Table S3. Bacteria were grown at 37° in Luria-Bertani (LB) agar plates or in 4 mL of liquid broth under vigorous shaking (250 rpm) unless indicated otherwise. Kanamycin (50 μg/ml) or Streptomycin (100 μg/mL for *E. coli* and 200 μg/ml for *Klebsiella*) were used in the selection of the plasmids strain selection and cultures. Antibiotics were purchased from Sigma.

### Growth media description

AUM and ASM stand for artificial urine medium and artificial sputum medium, respectively, and were done as described previously (Brooks and Keevil 1997, Fung et al 2010). Briefly, AUM is mainly composed of urea and peptone with trace amounts of lactic acid, uric acid, creatinine and peptone. ASM is composed of mucin, DNA, egg yolk and aminoacids. LB is composed of tryptone and yeast extract. M02 and M005 indicate minimal M63B1 supplemented with 0.2% and 0.05% glucose (sole carbon source), respectively.

### Determination of serotype

Complete or draft genomes were serotyped using Kaptive with default settings (Wick et al 2018).

### Natural loss of capsule

An overnight culture in LB was diluted at 1:100 in the different environments (ASM, LB, M02, M005, AUM) in a final volume of 4 mL. Cultures were allowed to grow for 24 hours at 37°C and diluted again to 1:100 in fresh media. This was repeated for 3 times. Then, each culture in each environment was serially diluted and CFUs were counted (3 plates per sample, > 100 colonies per plate). Non-capsulated mutants are easily visualized by the naked eye as mutants produce smaller, rough and translucent colonies. Results are expressed as a ratio of non-capsulated mutant colonies observed over the total plate count. The limit of detection of non-capsulated mutants in this assay was 0.22% ± 0.04 or 1:450 per biological replicate per environment. To ensure that the naturally-emerged non-capsulated colonies observed on plate are *bona fide* genetic mutants and not phase-variants, we randomly chose 10 populations, using the *sample* function in R. Three non-capsulated clones from each population were randomly selected and restreaked twice. All clones derived from the original non-capsulated colony remained non-capsulated, demonstrating that the loss of the capsule is genetic.

### Generation of *Δwza, ΔwcaJ andΔwbaP* capsule mutants

Isogenic capsule mutants were constructed by an in-frame deletion of *wza* by allelic exchange. Upstream and downstream sequences of each gene (> 500pb) were amplified using Phusion Master Mix (Thermo Scientific) then joined by overlap PCR. The resulting PCR product was purified using the QIAquick Gel Extraction Kit and then cloned with the Zero Blunt^®^ TOPO^®^ PCR Cloning Kit (Invitrogen) into competent *E. coli* DH5α strain. KmR colonies were isolated and checked by PCR. Cloned Zero Blunt^®^ TOPO^®^ plasmid was extracted using the QIAprep Spin Miniprep Kit, and digested for 2 hours at 37°C with ApaI and SpeI restriction enzymes and ligated with T4 DNA ligase (Promega) overnight at 16° to a double-digested pKNG101 plasmid. The ligation was transformed into competent *E. coli* DH5α pir strain, and again into *E. coli* MFD λ-pir strain (Ferrieres et al 2010), used as a donor strain for conjugation into *Klebsiella spp*. Conjugations were performed for 24 hours at 37°C. Single cross-over mutants (transconjugants) were selected on Streptomycin plates (200 µg/ml). To select for plasmid excision, transconjugants were grown for 16 hours at 25°C and double cross-over mutants were selected on LB without salt and supplemented with 5% sucrose at room temperature. To confirm the loss of the plasmid, colonies were tested for their sensitivity to streptomycin. From each double-recombination, a mutant and a wild type were isolated. All experiments were performed in parallel with the parental strain, deletion mutant and the associated wild type, to ensure that the phenotype observed was directly due to the deletion and not to other potential SNPs that could have accumulated in the genome during the mutant generation, and thus present in both the mutant and the wild type.

Deletion mutants were first verified by Sanger, and then checked for off-target mutations using Illumina. *ΔwbaP* and *ΔwcaJ* did not present any off-target mutations. Four *Δwza* strains presented off-target mutations. Strain #63 (KL107) *Δwza* has a 15bp deletion upstream of *rfaH*, a major capsule regulator. Inactivation of *rfaH* inhibits the initiation of capsule synthesis altogether. Strain #37 (KL24) has an extra 15 bp deletion just before the ATG of *wza*. Strain #51 (KL24) has a SNP in *pcaR*. There are no reports on a direct influence of this gene on capsule production. Strain #55 (KL2) has a SNP in *sacX* gene which results in delayed growth. Despite the fact that these strains are *bona fide* capsule mutants, as shown by microscopic observation and capsule quantification, the results are presented as supplemental figures.

### Microscopy

Bacteria capsules were visualized using Indian Ink coloration on an exponential phase culture. Images were acquired on a Nikon Eclipse E400 camera using the Digital Net DN100 camera control unit and a 100X objective.

### Capsule extraction and quantification

The bacterial capsule was extracted as described in (Domenico et al 1989) and quantified by using the uronic acid method (Blumenkrantz and Asboe-Hansen 1973). Briefly, 500 μL of an overnight culture was mixed with 100 μL of 1% Zwittergent 3-14 detergent in 100 mM citric acid (pH 2.0) and heated at 56°C for 20 minutes. After it was centrifuged for 5 min at 14,000 rpm and 300 μL of the supernatant was transferred to a new tube. Absolute ethanol was added to a final concentration of 80% and the tubes were placed on ice for 20 minutes. After a second wash with ethanol at 70%, the pellet was dried and dissolved in 250 μL of distilled water. The pellet was then incubated for 2 hours at 56°C. Polysaccharides were then quantified by measuring the amount of uronic acid. A 1,200 μL volume of 0.0125 M tetraborate in concentrated H_2_SO_4_ was added to 200 μL of the sample to be tested. The mixture was vigorously vortexed and heated in a boiling-water bath for 5 min. The mixture was allowed to cool, and 20 μL of 0.15% 3-hydroxydiphenol in 0.5% NaOH was added. The tubes were shaken, and 100 μL were transferred to a microtiter plate for absorbance measurements (520 nm). The uronic acid concentration in each sample was determined from a standard curve of glucuronic acid.

### Competition assay

Mutants and their respective wild types were grown overnight in LB. They were then mixed in a 1:1 proportion. A sample was taken and used for serial dilution and CFU counting as a control of T_0_. The co-culture was then diluted 1/100 in 4 mL of the different environments (i.e. LB, AUM, ASM, M02 and M005). For control experiments, ASM and LB were diluted 1:10 in M63B1 with no supplement of glucose. After 24 hours of competition (T_24_), each mixture was serially diluted and plated. Capsulated and non-capsulated colonies are clearly differentiated visually and counted separately. The competitive index of capsulated strains was calculated by dividing the ratio of CFU at T_24_ over T_0_.

### Growth curves

Overnight cultures were diluted at 1:100 in the aforementioned environments. 200 μl of each subculture was transferred in a 96-well microplate. Cell densities of cultures were estimated with a TECAN Genios™ plate reader. Absorbance (OD_600_). Light absorbance values of each well reflect the average of 6 reads per well. Absorbance values from within-block technical replicates were averaged and these averages were used as statistically independent data points. (i) *Growth rate*. Minimum generation times were estimated across replicates for the 1 hour interval (ΔT) spanning the fastest growth during exponential growth phase. This was calculated as:

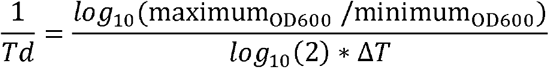

(ii) *Maximum yield*. Relative yields were calculated measuring ratios of the maximum absorbance of the assay compared to the reference (parental strain). The experiments were performed at least three independent times, with three technical replicates per experiment. The average of the three technical replicates per experiment was used for statistical purposes. Control experiments were performed to assess the correlation between CFU and OD_600_ during growth for both capsulated (Spearman’s rho = 1.0, P = 0.002) and non-capsulated mutants (Spearman’s rho = 0.94, P = 0.016). To confirm the differences in yield observed in minimal medium between capsulated and non-capsulated strains, we performed a control experiment (N = 3) in which we grew the strains #56 and #24 in microtiterplates as described before. After 16h, we plated an aliquot to count CFUs. Significantly less CFUs were observed in the non-capsulated mutants compared to wild type (P<0.05 for both strains). (iii) *Area under the curve* (AUC). AUC was calculated using the R function *trapz* from the *pracma* package.

### Serum resistance

The ability of *Klebsiella* strains to resist killing by serum was performed as described previously (Dorman et al 2018, Podschun et al 1993, Podschun and Ullmann 1998). Briefly, bacteria were grown in LB to OD 600 nm of 1, pelleted by centrifugation and resuspended in sterile PBS 1X. 200 μl of bacterial suspension was added to 400 μl of pre-warmed at 37°C human sera (Sigma-Aldrich S7023), and the mixture was incubated for 2h at 37°C. Control reactions were performed with serum inactivated by heat at 56°C for 30 min. Reactions were stopped by placing on ice, and viable bacterial counts were determined before and after incubation by serial dilution and CFU plating.

### Bile salts resistance

Bacteria were grown overnight in LB. As previously described (Tan et al 2020b), cultures were then serially diluted and plated in LB supplemented with either 0, 0.05% (physiological conditions) and 0.5% of CHO (primary bile salt, sodium cholate) or DCH (secondary bile salt, sodium deoxycholate). Colonies were allowed to grow and surviving CFU were counted after 24 hours. The average of two technical replicates per experiment was used for statistical purposes.

### Starvation

Bacteria were grown in LB to OD=0.2, 1mL of the culture was pelleted by centrifugation and, resuspended in 400 μl of PBS 1X, as in (Hamaguchi et al 2018). Cultures were allowed to starve at room temperature. Aliquots were taken at 0, 30 and 60 days, and used for serial dilutions and CFU plating.

### Biofilm Production

The ability of strains to form biofilm was performed as described in (Rendueles et al 2011). Overnight cultures were diluted at 1:100 in the different growth media mentioned earlier. 100 μL of each diluted culture was transferred in triplicate into 96-well microplate wells and allowed to grow for 24 hours without shaking at 37°C. Cells were gently removed from the wells and washed once in distilled water. To stain biofilms, 125 μL of 1% crystal violet was added to each well for 20 minutes. The crystal violet was gently decanted and each well was washed three times with distilled water. After the plate was totally dry, the biofilm was solubilized for 10 min in 150 μL of mix with 80% ethanol and 20% acetone. The absorbance of the sample was read at OD_570_. At least 3 biological independent replicates were performed for each sample.

### Hypermucoviscosity

The optical density OD_600_ of overnight cultures were first determined by spectrophotometry as described in (Dorman et al 2018). The cultures were then sedimented by slow centrifugation (at 2,500 X g) for 5 min at room temperature. The optical density OD_600_ of the top 650 μL of supernatant was determined. Results were expressed as a ratio of the supernatant OD_600_ to that of the input culture.

### Dessication test

Dessication assay was carried as previously described in (Chin et al 2018). Briefly, 25 μL of exponentially-growing *Klebsiella* strains were deposited at the bottom of microtiterplates, and allowed the culture to dry under a laminar flow hood. Strains were then left either at 37° for one, two or four days or at room temperature for two, four, seven and 14 days. PBS was added to resuspend the cultures and diluted serially from CFU counts.

### Quantification of dead cells

Cells were allowed to grow as described above with a minor modification; over day cultures were used to inoculate the 96-well microtiter plates. After 16 hours of growth in M02, dead cells were quantified using the LIVE/DEAD™ BacLight™ Bacterial Viability Kit from Invitrogen, following provider instructions. Images were acquired on a Nikon Eclipse E400 microscope using the Digital Net DN100 camera control unit and a 100X objective. Cells were counted using ImageJ software (https://imagej.net/). On average, 6 pictures were taken per sample (min = 3, max = 10) and *ca*. 550 cells per sample were observed (min = 161, max =1604). The experiment was performed four times.

### Growth on exogenously added capsule

Capsules were extracted from either wild type strains or their respective Δ*wcaJ* or Δ*wbaP* (for strain #24) mutants and quantified as described above. To initiate growth, overnight cultures of Δ*wcaJ* mutants were diluted at 1:100 in M63B1 media. Media was supplemented with either 0.8 ng/μL of extracted capsule from wild type or an equivalent volume extracted from the Δ*wcaJ* or Δ*wbaP* (for strain 24) mutants. As a negative control, cultures were allowed to grow in M63B1 with no nutrients. In parallel, M63B1 was supplemented with glucuronic acid (the standard used for estimating capsule concentration) at 0.8 ng/μL final. Growth was allowed for 48 hours at 37° and 250 rpm. Bacterial counts were performed at T_0,_ T_24_ and T_48_ by serial dilution and plating. Results are expressed as the increase in CFU over 24h.

## ACKNOWLEDGEMENTS

We thank Matthieu Haudiquet, Jean-Marc Ghigo and Nienke Buddelmeijer for critical reading of the manuscript. We thank Sylvain Brisse for providing us with the necessary Klebsiella strains, and Christiane Forestier and Damien Balestrino for providing the pKNG101 plasmid used to generate the mutants. We would also like to thank Jean-Marc Ghigo and Christophe Beloin for the gift of *E. coli* S17 MFD λ-pir strain and *E. coli* DH5α-pir.

## SUPPLEMENTAL FIGURE LEGENDS

**Figure S1. Capsule quantification of each strain in each growth media by the uronic acid method**.

**Figure S2. Ratio of capsulated colonies throughout 30 days of evolution in two media**. Strains were diluted 1:100 and inoculated in fresh media every 24 hours during 30 days. CFUs were counted every day. Each line represents an independent population (N = 3).

**Figure S3. Capsule quantification of each mutant by the uronic acid method**. This experiment was performed on overnight cultures grown in LB. Each dot corresponds to an independent experiment. The background noise of this assay is established around 1.5 ng/μL. Statistical tests: * P < 0.05; ** P < 0.01; *** P<0.001.

**Figure S4. Fitness advantage/disadvantage of capsulated strains in nutrient poor/rich media against *wza* mutants (A) or *wcaJ/wbaP* mutants (B)**. Competition between wild type and their isogenic capsule mutants in different growth media. Each point represents the average of at least three independent replicates. * P < 0.05 Tukey post hoc, p-values adjusted for multiple testing.

**Figure S5. Survival of strains under starvation and desiccation. A**. Survival of parental, mutant and wild type strains, relative to day 0. Data is log_10_ -transformed. Error bars are not displayed for clarity purposes. Colours represent different serotypes (as in Figure 1), whereas the shape represent different species. **B**. Cellular survival, in CFU, after 1, 2 or four days of desiccation at 37°C. Error bars for independent points are not shown for clarity purposes.

**Figure S6. Growth parameters of the strains used in this study. A**. Minimum generation time (**A**), maximum yield (**B**) and area under the growth curve (**C**) of the mutant and wild type strains, relative to the parental strain. Statistics were calculated using a paired t-test. * P <0.05, ** P < 0.01, *** P < 0.001. **D**. Percentage of dead cells after 16 hours growth in a microtiter plate well. Each symbol represents an independent experiment. No significant differences are found across genotypes.

**Figure S7. Log**_**10**_**-increase of CFU after 48 hours of growth**. Growth of *ΔwcaJ* (#26 and #56) and *ΔwbaP* (#24) mutants was followed for 24 and 48 hours. Cells were grown in M63B1 without nutrient source (Control), with 0.8 ng/μL of purified capsule (Cap+), an equivalent volume of extraction of a non-capsulated strain (Cap-), and 0.8 ng/μL of glucuronic acid (the standard used to quantify the capsule). Individual points represent four independent experiments. Relevant statistical comparisons are shown, (ANOVA with post-hoc Tukey correction for multiple comparisons)

**Figure S8. Resistance to bile salts at physiological concentrations (0**.**05%), and at 10X (0**.**5%)**. Results are presented as Log_10_ (relative to control, LB plates). * P < 0.05, ** P< 0.01, *** P < 0.001

**Figure S9. Biofilm formation of parental, mutant and wild type strains across growth media**. P values reflect paired t-tests. * P < 0.05, ** P< 0.01, *** P < 0.001

**Figure S10. Growth rate and maximum yield of each population in each environment**.

**Figure S11. Biofilm formation of all strains in all environments**.

**Figure S12**.**Mucoviscosity index of strains in all environments**.

**Figure S13. PCA analyses of capsule-related phenotypes**. Phenotypes are coloured by serotype. Each individual point represents a strain. PCA was performed using *prcomp* function in R and ggbiplot package for visualization purposes.

