## Supplementary material for "Selection for the bacterial capsule in the absence of biotic and abiotic aggressions depends on growth conditions": Mat Sup

### SUPPLEMENTAL FIGURES

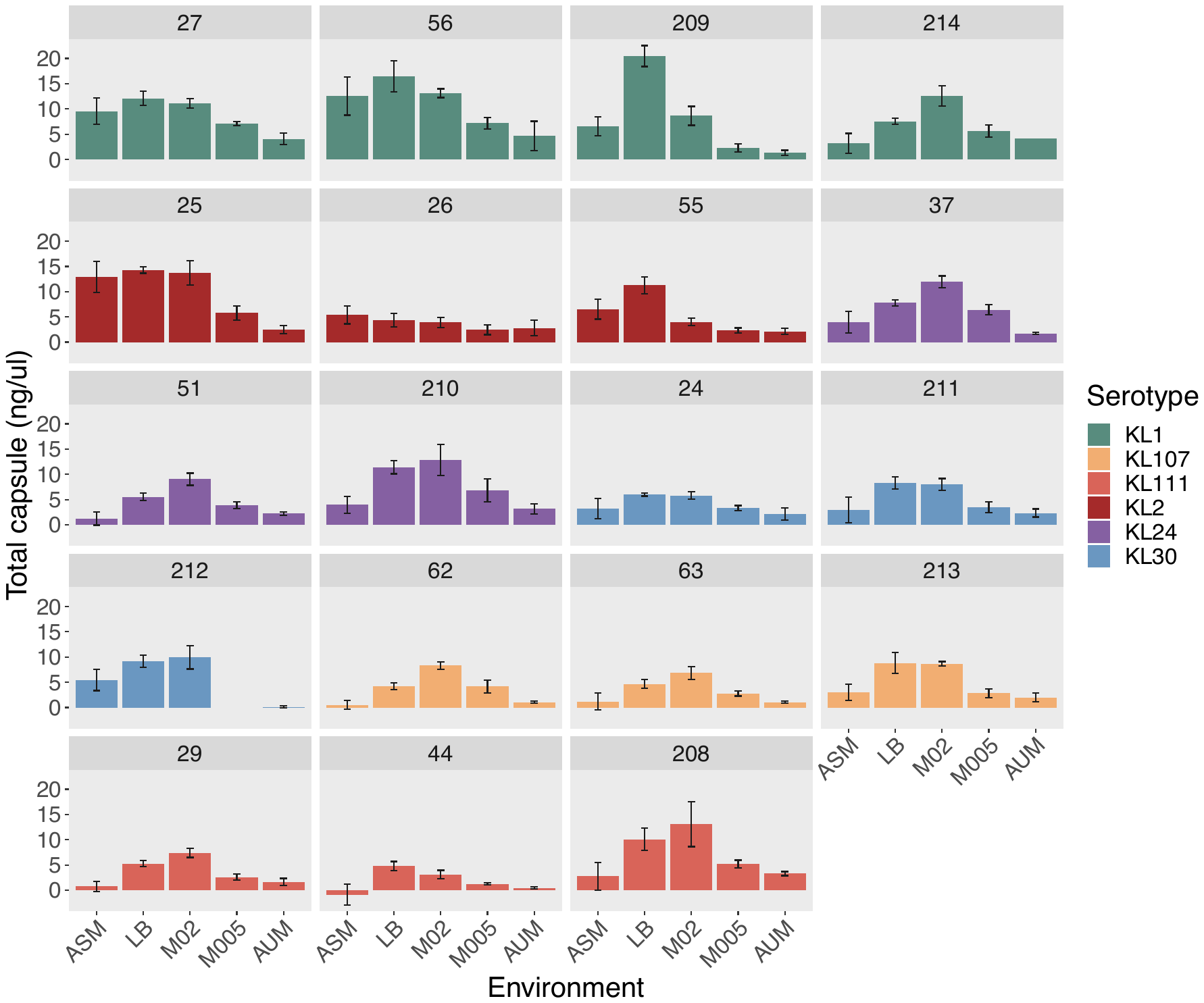

**Figure S1. Capsule quantification of each strain in each environment by the uronic acid**

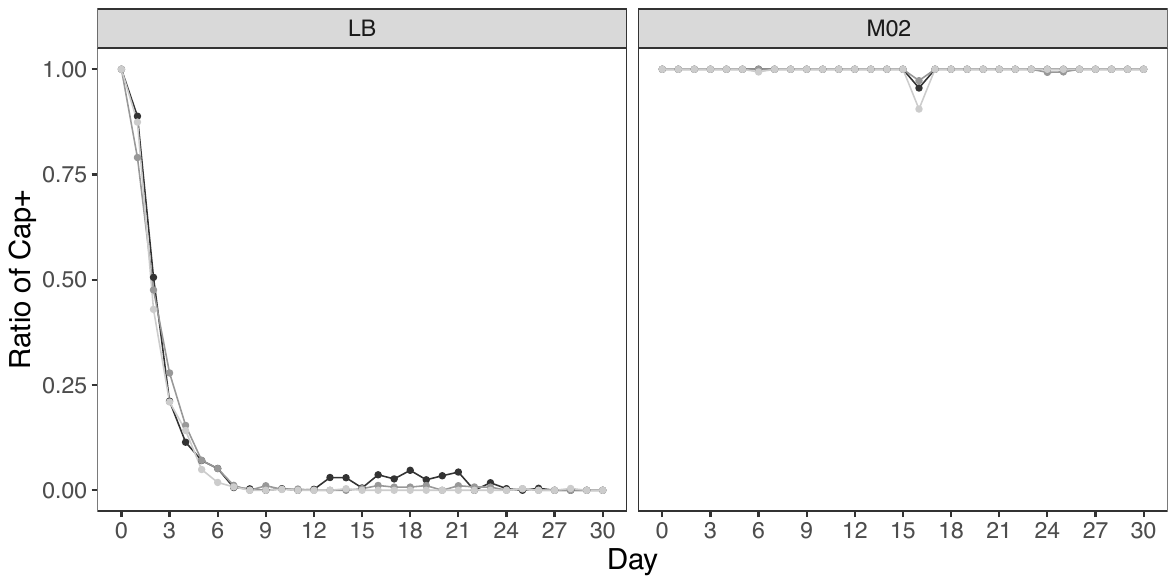

**Figure S2. Ratio of capsulated colonies throughout 30 days of evolution.** Strains were diluted 1:100 and inoculated in fresh media every 24 hours during 30 days. CFUs were counted every day. Each line represents an independent population (N = 3).

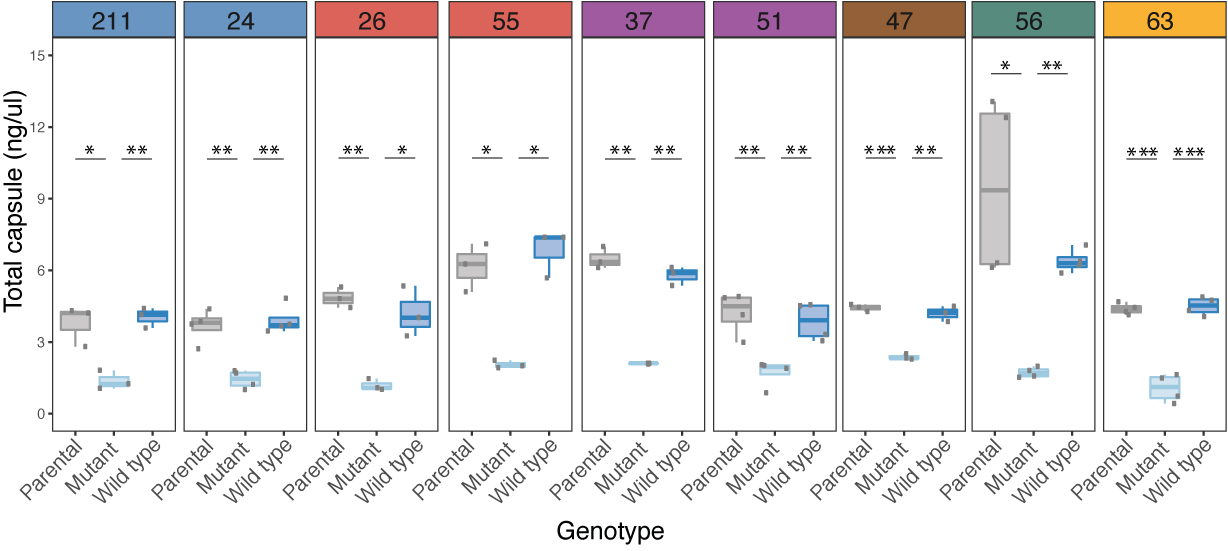

**Figure S3. Capsule quantification of each mutant by the uronic acid method**. This experiment was performed on overnight cultures grown in LB. Each dot corresponds to an independent experiment. The background noise of this assay is established around 1.5 ng/μL.

Statistical test: * P < 0.05; ** P < 0.01; *** P<0.001.

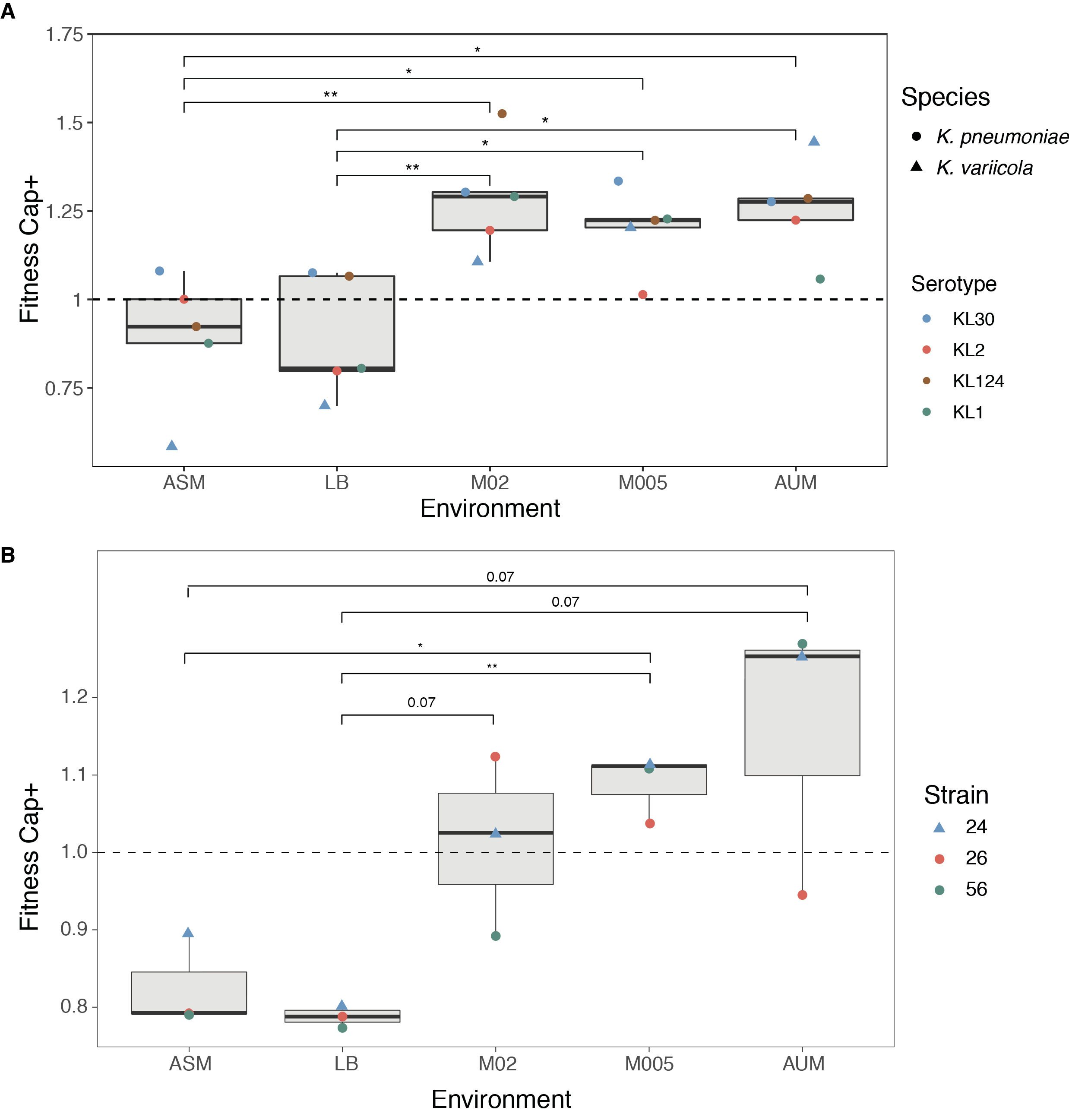

**Figure S4. Fitness advantage/disadvantage of capsulated strains in nutrient poor/rich media against *wza* mutants(A) or *wcaJ/wbaP* mutants (B).** Competition between wild type and their isogenic capsule mutants in different growth media. Each point represents the average of at least three independent replicates. * P < 0.05 Tukey post hoc, p-values adjusted for multiple testing.

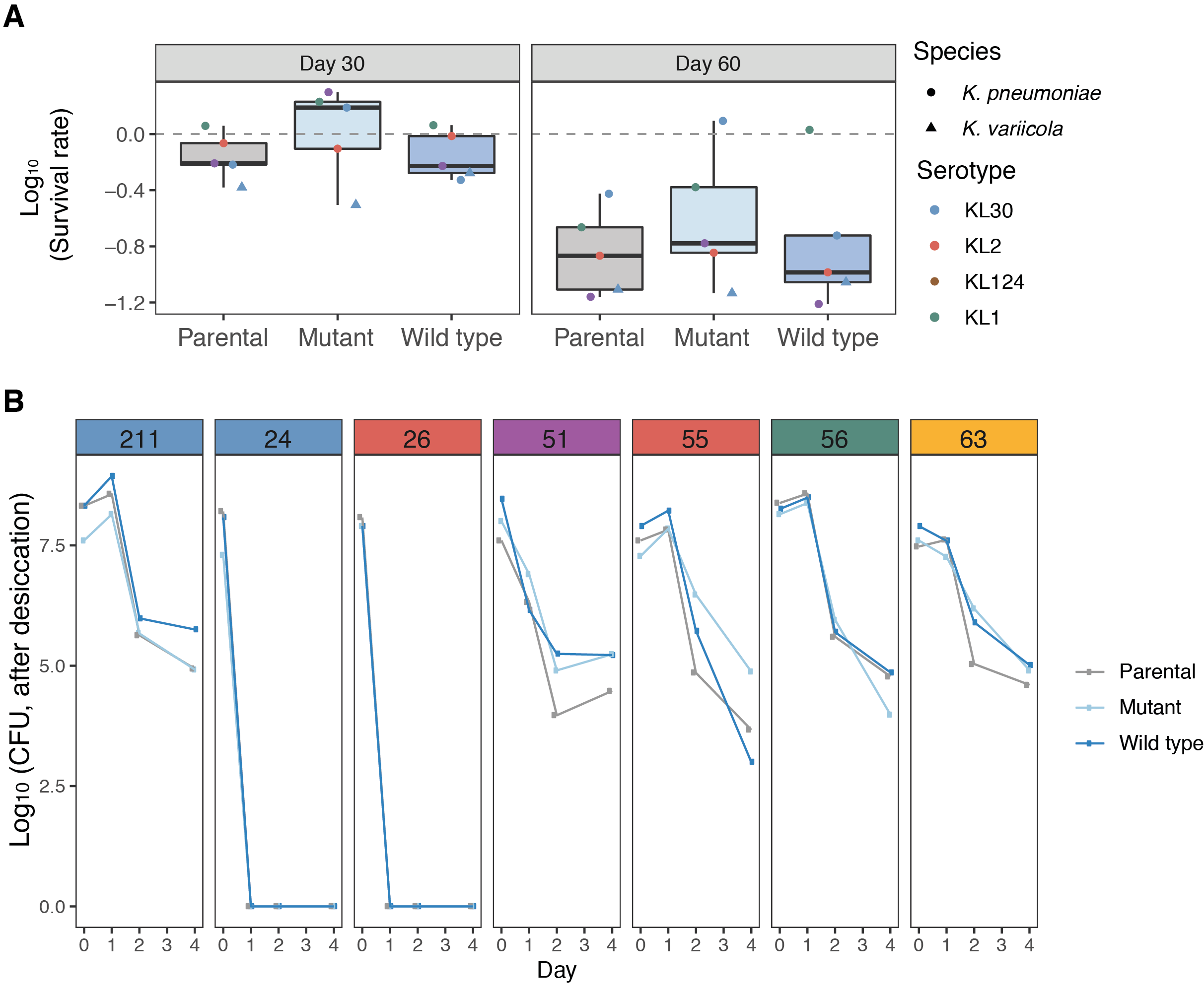

**Figure S5**. **Survival of strains under starvation and desiccation. A**. Survival of parental, mutant and wild type strains, relative to day 0. Data is log_10_ -transformed. Error bars are not displayed for clarity purposes. Colours represent different serotypes (as in Figure 1), whereas the shape represent different species. **B.** Cellular survival, in CFU, after 1, 2 or four days of desiccation at 37°. Error bars for independent points are not shown for clarity purposes.

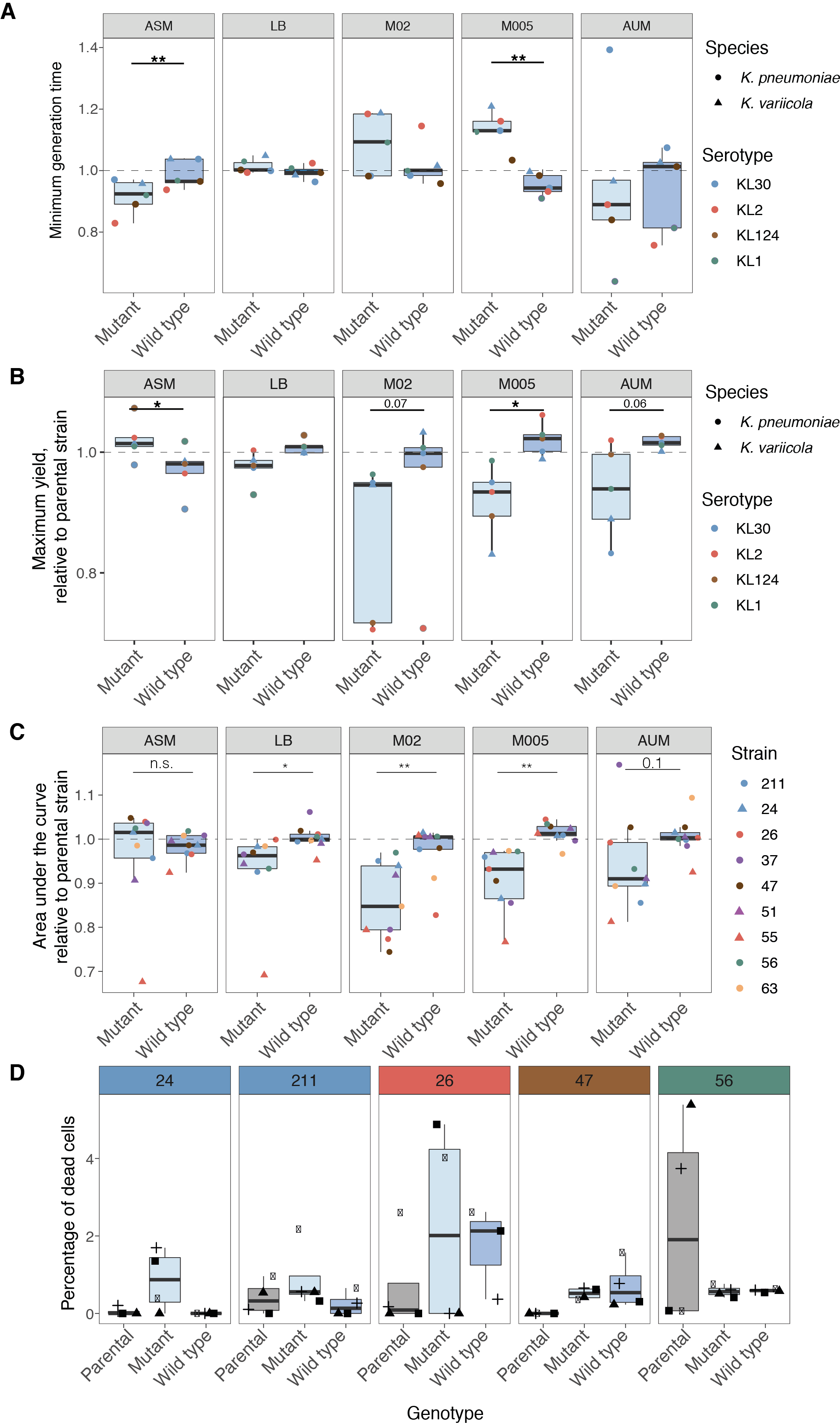

**Figure S6**. **Growth parameters of the strains used in this study. A.** Minimum generation time (**A**), maximum yield (**B**) and area under the growth curve (**C**) of the mutant and wild type strains, relative to the parental strain. Statistics were calculated using a paired t-test. * P <0.05,** P < 0.01, *** P < 0.001. **D.** Percentage of dead cells after 16 hours growth in a microtiter plate well. Each symbol represents an independent experiment. No significant differences are found across genotypes.

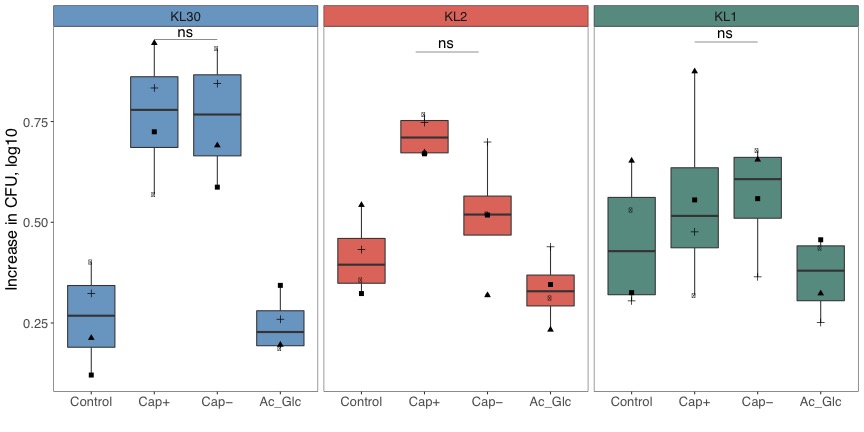

**Figure S7**. **Log_10_-increase of CFU after 48 hours of growth.** Growth of *ΔwcaJ* (#26 and #56) and *ΔwbaP* (#24) mutants was followed for 24 and 48 hours. Cells were grown in M63B1 without nutrient source (Control), with 0.8 ng/μL of purified capsule (Cap+), an equivalent volume of extraction of a non-capsulated strain (Cap-), and 0.8 ng/μL of glucuronic acid (the standard used to quantify the capsule). Individual points represent four independent experiments. Relevant statistical comparisons are shown, (ANOVA with post-hoc Tukey correction for multiple comparisons)

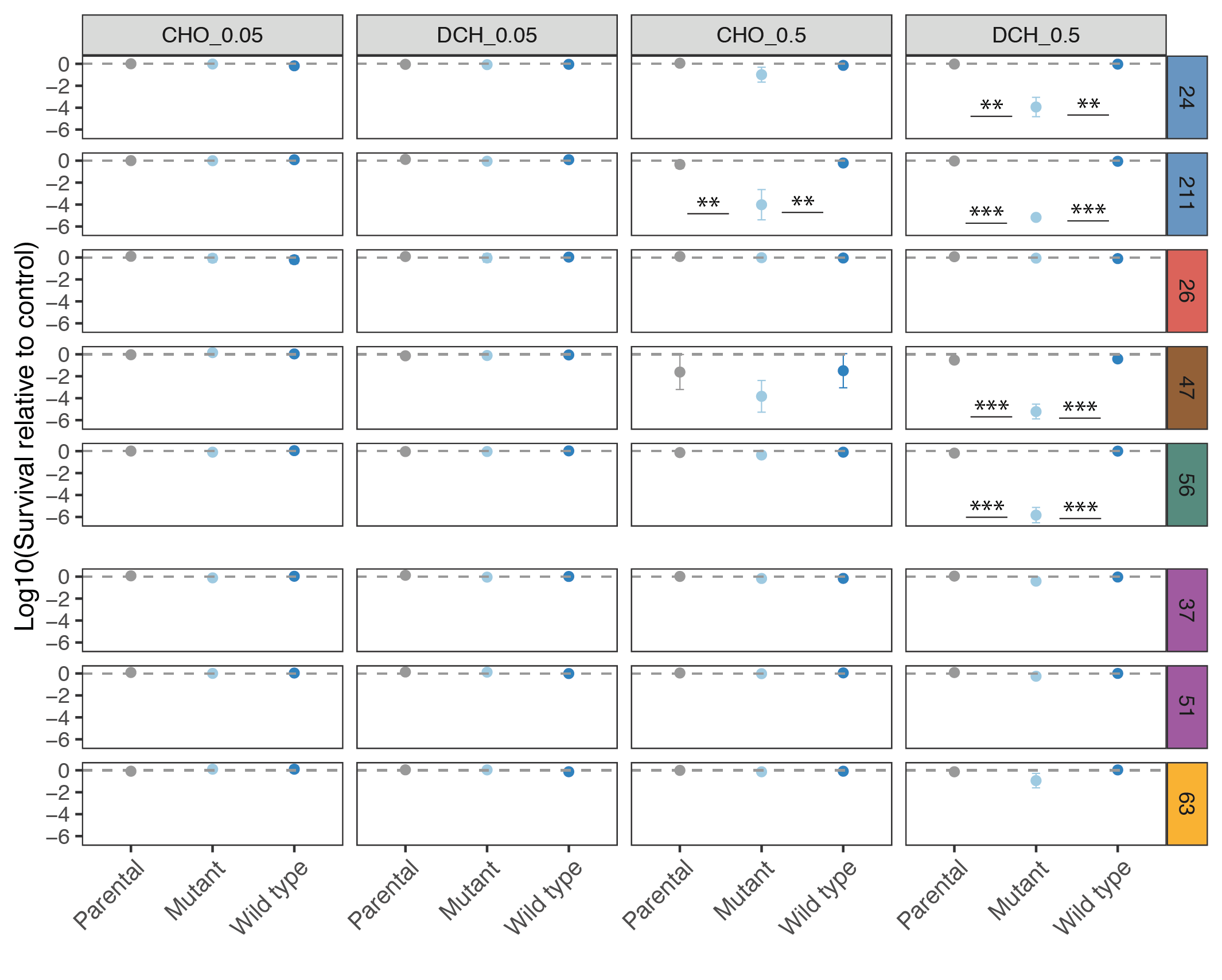

**Figure S8.** **Resistance to bile salts at physiological concentrations (0.05%), and at 10X (0.5%).** Results are presented as Log_10_ (relative to control, LB plates). * P < 0.05, ** P< 0.01, *** P < 0.001

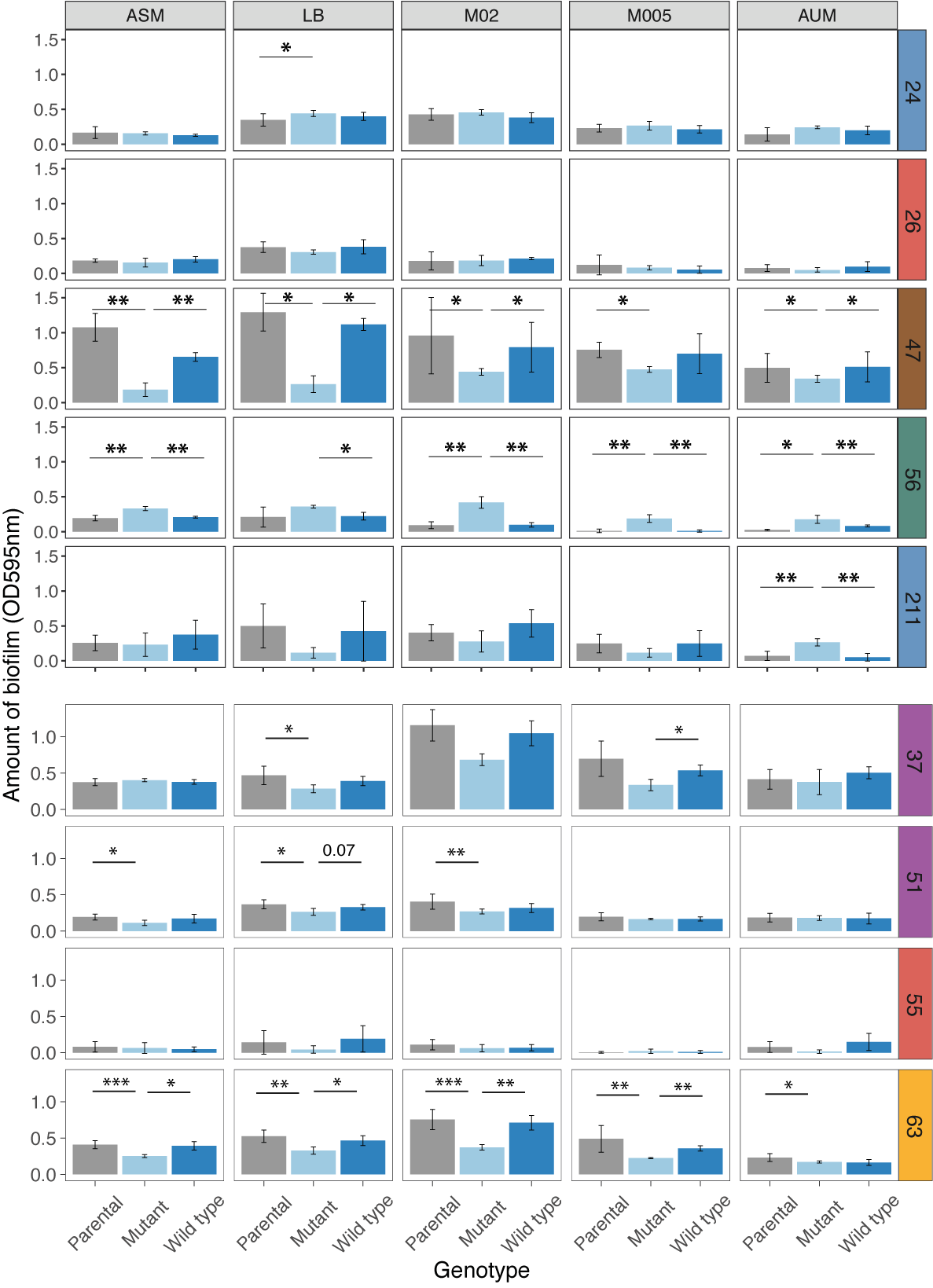

**Figure S9. Biofilm formation of parental, mutant and wild type strains across growth media.** P values reflect paired t-tests. * P < 0.05, ** P< 0.01, *** P < 0.001

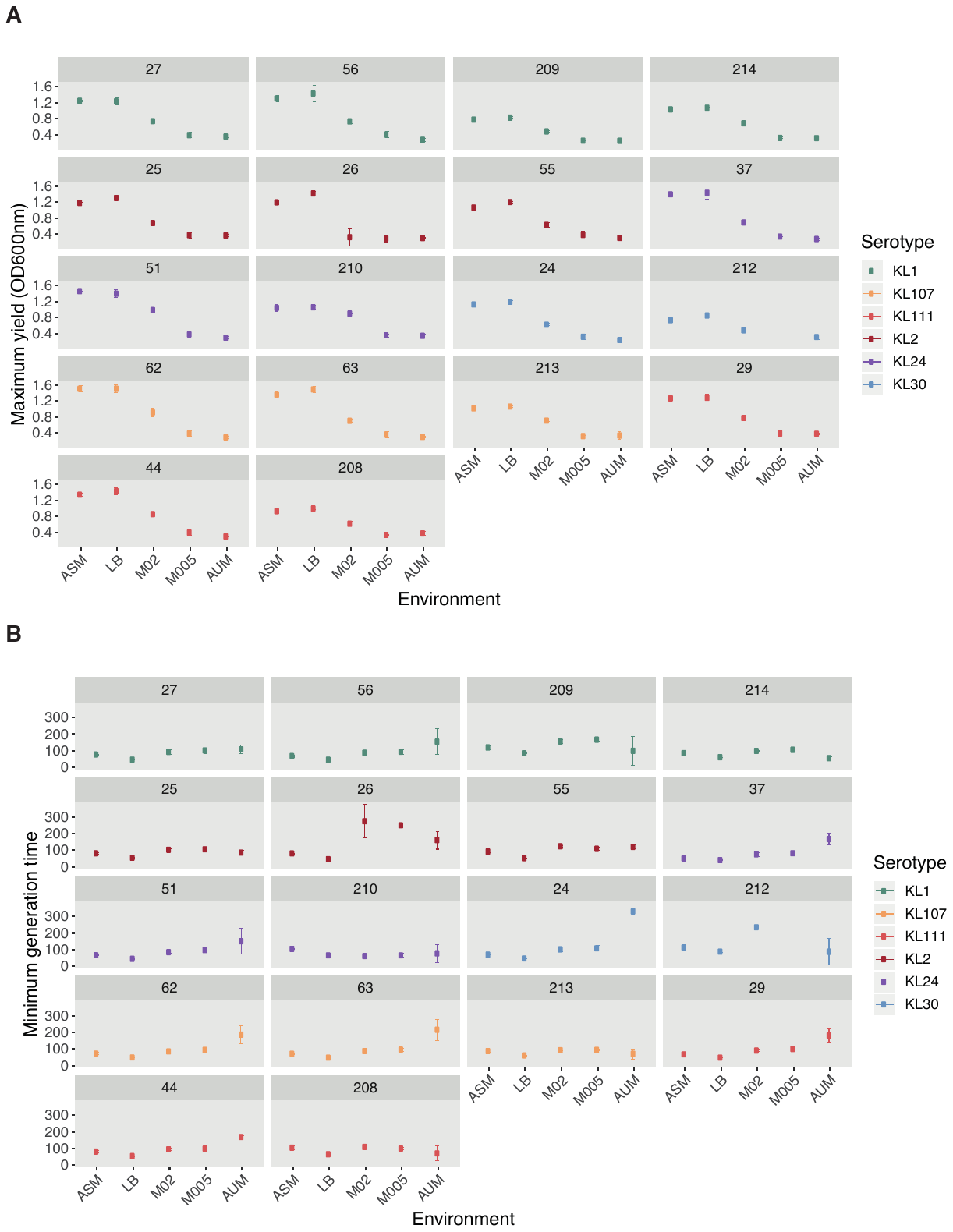

**Figure S10. Growth rate and maximum yield of each population in each environment.**

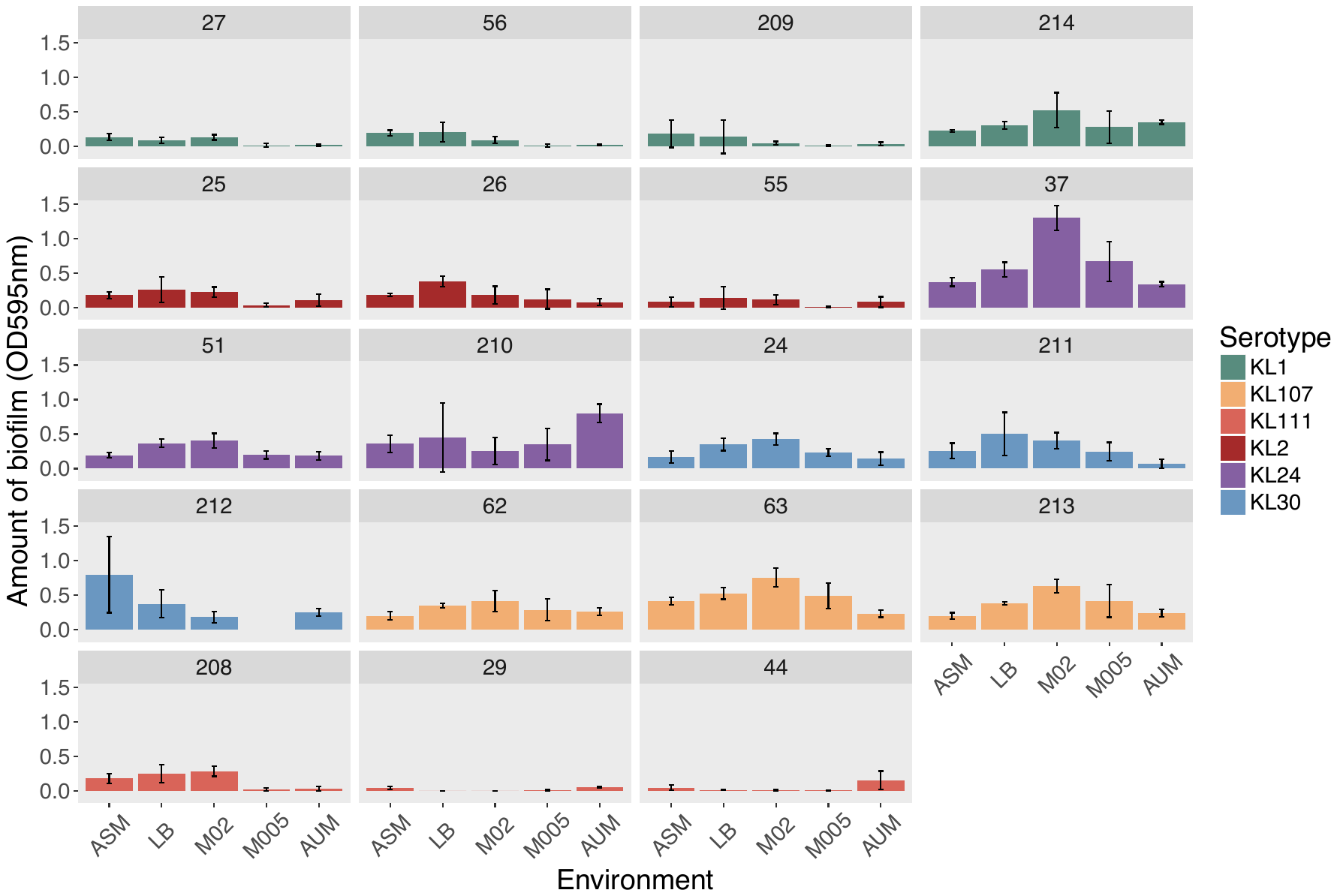

**Figure S11. Biofilm formation of all strains in all environments.**

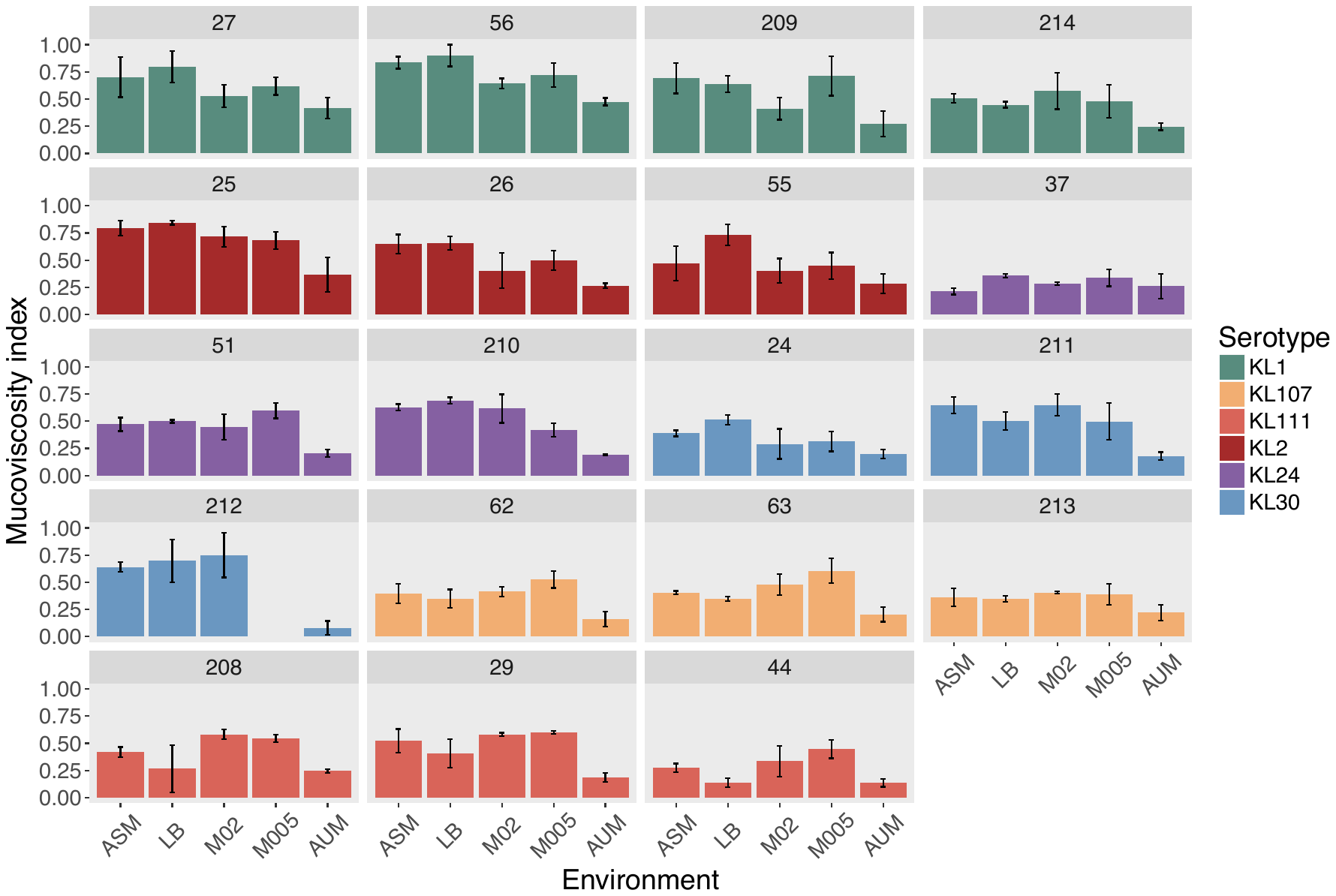

**Figure S12.Mucoviscosity index of strains in all environments.**

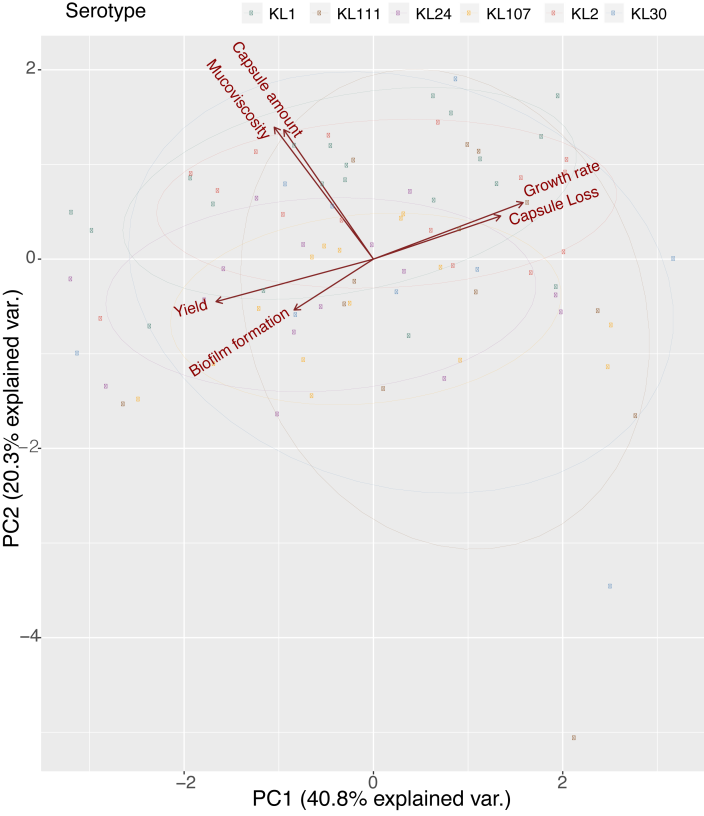

**Figure S13.PCA analyses of capsule-related phenotypes.** Phenotypes are coloured by serotype. Each individual point represents a strain. PCA was performed using *prcomp* function in R and ggbiplot package for visualization purposes.

### SUPPLEMENTAL TABLES

**Table S1. Strains and plasmids used in this study.**

| **# of strain** | **Strain name** | **ST** | | | **Species** | **Capsule serotype** | **O- Antigen serotype** | | | **Country** | | | **Isolation** | **Reference** |
| --- | --- | --- | --- | --- | --- | --- | --- | --- | --- | --- | --- | --- | --- | --- |
| *Klebsiella spp.* | | | | | | | | | | | | | | |
| 27 | SA12 | ST23 | | | *K. pneumoniae* | KL1 | O1v2 | | | France | | | Liver abcess | (1) |
| 56 | NTHU K2044 | ST23 | | | *K. pneumoniae* | KL1 | O1v2 | | | Taiwan | | | Liver abcess | (2) |
| 209 | CIP 52.208 | ND | | | *K. pneumoniae* | KL1 | O1v2 | | |  | | |  | (3) |
| 214 | ST254 | ST254 | | | *K. quasipneumoniae subsp similipneumoniae* | KL1 | O3/O3a | | | Cambodia | | |  | (1) |
| 25 | 03-9138 | ST65 | | | *K. pneumoniae* | KL2 | O1v2 | | | Martinique | | | Anal abcess | (4) |
| 26 | BJ1 | ST380 | | | *K. pneumoniae* | KL2 | O1v1 | | | France | | | Liver abcess | (5) |
| 55 | CIP 52.145 | ST66 | | | *K. pneumoniae* | KL2 | O1v2 | | | Indonesia | | | Clinical strain | (6) |
| 37 | 04A025 | ST15 | | | *K. pneumoniae* | KL24 | O1v1 | | | France | | | Blood, MDR | (7) |
| 51 | SB 1170 | ST45 | | | *K. pneumoniae* | KL24 | O2v1 | | | Netherlands | | | Carriage, feces | (7) |
| 210 | ST59 | ST59 | | | *K. pneumoniae* | KL24 | O1v1 | | |  | | |  | (3) |
| 24 | 342 | ST146 | | | *K. variicola* | KL30 | O3/O3a | | | USA | | | Environmental, corn | (8) |
| 211 | ST43 | ST43 | | | *K. pneumoniae* | KL30 | O1v1 | | |  | | |  | (1) |
| 212 | ST2435 | ST2435 | | | *K. pneumoniae* | KL30 | O1v1 | | |  | | |  | (1) |
| 62 | Kp13 | ST442 | | | *K. pneumoniae* | KL107 | O2v2 | | | Brazil | | | Clinical, MDR | (9) |
| 63 | NJST258-1 | ST258 | | | *K. pneumoniae* | KL107 | O2v2 | | | USA | | | Clinical, MDR | (10) |
| 213 | ST16 | ST16 | | | *K. pneumoniae* | KL107 | O2v2 | | | Cambodia | | |  | (1) |
| 208 | O81082 | ST20 | | | *K. pneumoniae* | KL111 | O3b | | |  | | |  | (1) |
| 29 | SB610 | ST2276 | | | *K. quasipneumoniae subsp similipneumoniae* | KL111 | O1v1 | | | Netherlands | | | Environmental, water | (7) |
| 44 | SB611 | ST63 | | | *K. pneumoniae* | KL111 | ND | | | Netherlands | | | Environmental, water | (7) |
| 47 | SB 617 | ST55 | | | *K. pneumoniae* | KL124 | ND | | | Netherlands | | | Environmental, water | (7) |
| Strains used for mutant construction | | | | | | | | | | | | | | |
| 100 | *E. coli S17*MFD  λ pir | |  | | *E. coli* |  | |  | | | laboratory strain | | | (11) |
| 287 | *E. coli* DH5⍺ pir | | | | *E. coli* |  |  | | | laboratory strain | | | |  |
| Plasmids used for mutant construction | | | | | | | | | | | | | | |
|  | pKNG101 |  | | | *(E. coli)* |  |  | | |  | | |  | (12) |
|  | pCR Blunt (TOPO) | | |  | *(E. coli)* |  | | |  | | |  | |  |

**Table S2. Multifactorial ANOVA Analyses.** R^2^ were calculated as 1 – deviance/null deviance from a general linear model, using the R function *glm*.

| **Y** | **Variable** | **F value** | **Pr(>F)** | **R^2^** |
| --- | --- | --- | --- | --- |
| Capsule amount | Environment | 21.41 | *< 0.0001* | 0.69 |
|  | Serotype | 6.21 | *0.0001* |  |
|  | Environment * Serotype | 0.95 | 0.86 |  |
| Biofilm | Environment | 2.38 | 0.06 | 0.57 |
|  | Serotype | 11.58 | *< 0.0001* |  |
|  | Environment * Serotype | 0.64 | 0.52 |  |
| Capsule loss | Environment | 13.15 | *< 0.0001* | 0.52 |
|  | Serotype | 0.86 | 0.51 |  |
|  | Environment * Serotype | 0.35 | 0.99 |  |
| Yield | Environment | 141.77 | *< 0.0001* | 0.90 |
|  | Serotype | 3.12 | 0.014 |  |
|  | Environment * Serotype | 0.59 | 0.89 |  |
| Growth rate | Environment | 8.80 | *< 0.0001* | 0.51 |
|  | Serotype | 2.81 | 0.02 |  |
|  | Environment * Serotype | 0.72 | 0.79 |  |
| Mucoviscosity | Environment | 14.8 | *< 0.0001* | 0.65 |
|  | Serotype | 6.7 | *< 0.0001* |  |
|  | Environment * Serotype | 0.97 | 0.51 |  |

**Table S3. Primers used in this study**

| **Primer name** | | **Sequence** | **Sense** | **Construction** |
| --- | --- | --- | --- | --- |
| **Construction of *wza*mutants** | | | | |
| KL124.wza.500-3 | GCATCGGCCTGATAAATTGG | | Reverse | capsule wza mutant strain # 47 |
| KL124.wza.500-5 | GCTTCTACGGACAGATGATCG | | Forward | capsule wza mutant strain # 47 |
| KL124.wza.atg3 | AATGTCACATCATCAGTAAATC | | Reverse | capsule wza mutant strain # 47 |
| KL124.wza.L5 | GACAGCTGCTCAGGAATTGGCAAATTTTGATTTACTGATGATGTGACATTTTATGTTTAATTCTATCCTAGTTGTTTGTTATGTTTAATTCTATCCTAGTTG | | Forward | capsule wza mutant strain # 47 |
| KL124.wza.out-3 | GTAATTTCTGGTGCTGACTGAGG | | Reverse | check PCR capsule wza mutant strain # 47 |
| KL124.wza.out-5 | ATACCGGGACAGATAATGAACC | | Forward | check PCR capsule wza mutant strain # 47 |
| KL24.wza.500-5 | GGGTAAGGATGCAGTGAACTGG | | Forward | capsule wza mutant strain # 51& #37 |
| KL24.wza.500-3 | CGATAAGCTCACCAACGATACG | | Reverse | capsule wza mutant strain # 51 & #37 |
| KL24.wza.L5 | TAACACAGCTGCTCAGGAATTGGCAAATTTTGGTTTACTGATGATGTGACATTTAatgtttgattcaatattagtaatttgtactgg | | Forward | capsule wza mutant strain # 51 & #37 |
| KL24.wza.atg-3 | AATGTCACATCATCAGTAAACC | | Reverse | capsule wza mutant strain # 51 & #37 |
| KL24.wza.out-5 | AGCCAGATGAATCAATATACCG | | Forward | check PCR capsule wza mutant strain # 51& #37 |
| KL24.wza.out-3 | CGCATCTTGTGCTGCTTGTCG | | Reverse | check PCR capsule wza mutant strain # 51& #37 |
| KL107.wza.500-5 | AAGCACATGATACGCGCACC | | Forward | capsule wza mutant strain # 63 |
| KL107.wza.500-3 | ATCAATACCTTCTGAGTCTCG | | Reverse | capsule wza mutant strain # 63 |
| KL107.wza.L5 | GGGAAGCAACGGCTATGACAGCTGCTCAGGAATTGGCAAATTTTGATTTACTGATGATGTGACATTTCatgtttaactcgattctagtagtttg | | Forward | capsule wza mutant strain # 63 |
| KL107.wza.atg-3 | AATGTCACATCATCAGTAAATC | | Reverse | capsule wza mutant strain # 63 |
| KL107.wza.out-5 | CAATATACCGCTGTGCCTCATG | | Forward | check PCR capsule wza mutant strain # 63 |
| KL107.wza.out-3 | CAACTCGGGCGTTTCATTATCG | | Reverse | check PCR capsule wza mutant strain # 63 |
| 25.wza.L3 | gcgaacggcatatattccctgtgcaaacaattaatattgtactaaacatAATGTCACATCATCAGTAAATCAAAATTTGC | | Reverse | capsule wza mutant strain # 55 |
| 25.wza.L5 | ATGTTTAGTACAATATTAATTGTTTGC | | Forward | capsule wza mutant strain # 55 |
| 25.wza-500-3 | TCACCAACCATTCGACCAAAG | | Reverse | capsulewza mutant strain # 55 reverse |
| SB3332.wza.out-5 | ACCGGGACAGATAACGAACC | | Forward | check PCR capsule wza mutant strain # 55 |
| SB3332.wza.out-3 | CACTTAACCTTGCCCATCCACG | | Reverse | check PCR capsule wza mutant strain # 55 |
| 25.wza-500-5 | TTAACTGGTATGTGGAAGCGCATG | | Forward | capsule wza mutant strain # 55 |
| **Verification of intermediate constructions** | | | | |
| M13_F | GTAAAACGACGGCCAG | | Forward | PCR check for insert in pCR Blunt (TOPO) |
| M13_R | CAGGAAACAGCTATGAC | | Reverse | PCR check for insert in pCR Blunt (TOPO) |
| pKNG101.verif3_SpeIout | ACCAAGCCTATGCCTACAGC | | Reverse | PCR check for insert in pKNG101 |
| pKNG101.verif5 | CTACATATCACAACGTGCGTGG | | Forward | PCR check for insert in pKNG101 |

| **Construction of *wbaP/wcaJ* mutants** | | | |
| --- | --- | --- | --- |
| 24. wbaP.500-3 | GATACATGGCTCACCGTCTTCTG | Reverse | capsule wbaP mutant strain #24 |
| 24. wbaP.out-3 | CGTTCCATTCGGTGAAGGTTTGC | Reverse | check PCR capsule wbaP mutant strain #24 |
| 24. wbaP.L5 | GTGAATATTTATAAATATCTTAATGGATTAGTGATTCTAAACAGTTCAGAATTTACAAAAGAGTAAATAACTatttgtaatgcgcttaaagtttgctctg | Forward | capsule wbaP mutant strain #24 |
| 24. wbaP.Atg-3 | agttatttactcttttgtaaattc | Reverse | capsule wbaP mutant strain #24 |
| 24. wbaP.500-5 | actatcgggattagttttctcg | Forward | capsule wbaP mutant strain #24 |
| 24. wbaP.out-5 | ggagtattatacaaggcctgaccg | Forward | check PCR capsule wbaP mutant strain #24 |
| KL2.wcaJ.L5 | gttacgttaatgaaacttatcgacagatatgcttgtttataaatgagtgattgaattctagatgctccttaagacaagg | Forward | capsule wcaJ mutant strain #26 |
| KL2.wcaJ.500-3 | CACCATACTCAATGCCGTTATGC | Reverse | capsule wcaJ mutant strain #26 |
| KL2.wcaJ.L3 | aattcaatcactcatttataaac | Reverse | capsule wcaJ mutant strain #26 |
| KL2.wcaJ.500-5 | ttcttaaacttaagaacataagagc | Forward | capsule wcaJ mutant strain #26 |
| KL2.wcaJ.out-5 | GAATGGAATTGTTCTGCCTCTGAC | Forward | check PCR capsule wcaJ mutant strain #26 |
| KL2.wcaJ.out-3 | CTCAGTTCACCTTCGTTCCATTCG | Reverse | check PCR capsule wcaJ mutant strain #26 |
| 56.wcaJ_2.600-5 | GCTCGGACCTGAATTTGGATAATGTC | Forward | capsule wcaJ mutant strain #56 |
| 56.wcaJ_2.600-3 | CTCATAGGCTTCTTTCTGCCCACC | Reverse | capsule wcaJ mutant strain #56 |
| 56.wcaJ_2.STOP-5 | TAAGATGCTGCTTAAGATAAGC | Forward | capsule wcaJ mutant strain #56 |
| 56.wcaJ_2.L3 | GTTAACTAGCACAACATATAAATTTTGGACTATAAATGCACAATGCTTATCTTAAGCAGCATCTTAGAAGATTTCCTTAAAATATTGAACC | Reverse | capsule wcaJ mutant strain #56 |
| KL1.wcaJ.out5 | AGATTTCGAGGTCGATGCTATGC | Forward | check PCR capsule wcaJ mutant strain #56 |
| KL1.wcaJ.out3 | TCGGTTATCAGAGACAGAGGTTC | Reverse | check PCR capsule wcaJ mutant strain#56 |
